# Type I interferon dynamics determines memory B cell epigenetic identity in chronic viral infection

**DOI:** 10.1101/2023.03.02.530850

**Authors:** Lucy Cooper, Hui Xu, Jack Polmear, Christopher Szeto, Ee Shan Pang, Mansi Gupta, Justin J Taylor, Katherine J L Jackson, Angela Nguyen, Nicole La Gruta, Luciano Martelotto, Ian A Parish, Meredith O’Keeffe, Christopher D Scharer, Stephanie Gras, Kim L Good-Jacobson

**Affiliations:** Department of Biochemistry and Molecular Biology, Monash University, Clayton, Victoria, Australia; Immunity Program, Biomedicine Discovery Institute, Monash University, Clayton, Victoria, Australia; Department of Biochemistry and Chemistry, La Trobe Institute for Molecular Science, La Trobe University, Melbourne, Victoria, Australia; Department of Microbiology and Immunology, School of Medicine, Emory University, Atlanta, GA, USA; Vaccine and Infectious Disease Division, Fred Hutchinson Cancer Center; Garvan Institute of Medical Research, Darlinghurst, NSW, Australia; Adelaide Centre for Epigenetics and the South Australian Immunogenomics Cancer Institute, Faculty of Health and Medical Sciences, The University of Adelaide, Adelaide, SA, Australia; University of Melbourne Centre for Cancer Research, Victoria Comprehensive Cancer Centre, Melbourne, VIC, Australia; Peter MacCallum Cancer Centre, Melbourne, Victoria, Australia; Sir Peter MacCallum Department of Oncology, The University of Melbourne, Parkville, Victoria, Australia; John Curtin School of Medical Research, ANU, ACT, Australia

**Keywords:** B cells, chronic viral infection, IFN, epigenetics, memory B cells, LCMV, atypical, long covid, scATAC-sequencing

## Abstract

Memory B cells are key providers of long-lived immunity against infectious disease, yet in chronic viral infection they do not produce effective protection. How chronic viral infection disrupts memory B cell development, and whether such changes are reversible, remains unknown. Here, we uncover type-I interferon (IFN-I) dynamics as a key determinant in shaping chronic memory B cell development. Through single-cell (sc)ATAC-sequencing and scRNA-sequencing, we identified a unique memory subset enriched for IFN-stimulated genes (ISGs) during chronic lymphocytic choriomeningitis virus infection. Blockade of IFNAR-1 early in infection transformed the chromatin landscape of chronic memory B cells, decreasing accessibility at ISG-inducing transcription factor binding motifs and inducing a phenotypic change in the dominating memory B cell subset. However, timing was critical, with memory B cells resistant to intervention after 4 weeks post-infection. Together, our research identifies a key mechanism to instruct memory B cell identity during viral infection.

**One Sentence Summary:** IFN dynamics in chronic versus acute viral infection determines memory B cell development.

## INTRODUCTION

A fundamental pillar of immunity are immune memory populations, antigen-experienced cells with increased capabilities to persist and rapidly reactivate upon reinfection. Memory B cells drive superior antibody-mediated responses upon re-infection compared to the primary response^1–3^, enabling the rapid clearance of an infection before it can cause severe disease. However, chronic viral infections such as HIV, hepatitis C, cytomegalovirus and potentially SARS-CoV-2 can disrupt memory B cell development and antibody production, leading to incomplete and ineffective immune protection^4, 5^. The key molecular and microenvironmental determinants of chronic memory B cell development are still mostly unknown, thus limiting our ability to therapeutically redirect memory B cell development to functional subsets.

The microenvironment plays a crucial role in shaping the immune response against viral infections. The interplay between viral replication and inflammatory mediators, such as interferons, not only charts the course for whether an infection will be cleared effectively but also whether the immune response becomes dysfunctional^6–9^. To understand how humoral memory is disrupted by chronic infection, much research has focused on understanding the shift in the phenotype of memory B cells that is common across several severe infectious and inflammatory diseases^5, 10^. Antigen-experienced cells that have downregulated CD21 and CD27 and may upregulate FCRL5, T-bet and/or CD11c are detected at elevated frequency in the peripheral blood of HIV, hepatitis C, cytomegalovirus, systemic lupus erythematosus and Long COVID patients^5, 9, 11–16^. Cells of this phenotype can also be generated during acute viral infection^17, 18^, and thus it is still unclear how chronic viral infection rewires memory B cell population to cause dysfunction, and why these changes may not be completely reverted by treatment. Antiretroviral therapy in HIV patients, for example, restores the prominence of the conventional memory B cell phenotype, but if viral loads rebound, memory B cell subsets revert back to a disease state^19^. We do not know when, or if, the point of no return for recovery of memory B cell function occurs in chronic viral infection^20^. It is therefore vital that we better understand the fundamental changes to memory B cells that continual pathogenic pressure imparts.

During an immune response, antigen-activated B cells react to the signals within their pathogen-induced microenvironment and enact large-scale changes to their gene expression program to induce differentiation into antibody-secreting cells, germinal center (GC) B cells or memory B cells. Epigenetic regulation facilitates the integration of these signals by enabling new gene expression programs^4, 21, 22^. As a result of chronic viral pathogens and/or chronic antigen exposure, memory T cells become functionally exhausted, which has been linked to changes in the epigenome^21, 23, 24^. In contrast, little is known about (i) how memory B cells are fundamentally altered by sustained epigenetic changes during chronic infection, (ii) whether there are key microenvironmental differences between acute and chronic infection that determine the type of memory B cell produced, and (iii) if chronic memory B cells can be epigenetically rewired to resemble conventional memory B cells upon administration of therapeutics or whether they are resistant to recovery. Thus, improved resolution of the molecular architecture and regulation of distinct memory B cell subsets that arise during an immune response, particularly in chronic disease, is key to enabling therapeutic targeting to drive effective humoral memory.

We set out to determine how and when the microenvironment during chronic viral infection codifies intrinsic changes within memory B cells. We established a system in which memory B cells formed in chronic infection, compared to acute infection of the same virus, mimicked those in patients with ongoing severe viral infection. Single-cell ATAC-sequencing (scATAC-seq) and scRNA-seq defined the discrete subsets that arose during chronic or acute lymphocytic choriomeningitis virus (LCMV) infection. We identify that memory B cells which significantly expand in the spleen during chronic infection were enriched for IFN-stimulated genes (ISGs) and were independent of the T-bet^+^ subset normally associated with chronic infection. The chronic memory B cell chromatin landscape was amenable to remodelling but relied on the timing of therapeutic intervention. Blocking type I IFN signaling early in the response both increased cell numbers and changed the ascendant epigenetic signature of memory B cells. In contrast, administration of IFN blocking antibodies at 4 weeks post-infection or modulation of viral load did not convert the profile of chronic memory B cells, suggesting that dysfunctional imprinting of chronic memory B cells occurs early in the response, and was not due solely to persistent viral load. Collectively, our study reveals the consequences of delayed IFN induction and magnitude on memory B cell identity in chronic infection.

## RESULTS

### Formation of antigen-specific memory B cells during a polyclonal response to chronic versus acute LCMV infection

We set out to investigate how the memory B cell epigenetic landscape is remodelled by a chronic viral infection-induced microenvironment. To do this, we established a system in which memory B cell formation was assessable during a polyclonal response to acute versus chronic viral infection. We used the comparative LCMV model, with which we could study B cell differentiation in response to two strains of the same virus: one that induces an acute infection resolved by two weeks (LCMV-WE), or a chronic, persisting infection (LCMV-Docile)^4, 25, 26^. The strength of this model is that B cell-differentiation in response to the same virus can be tracked and assessed in either an acute or chronic setting, overcoming potential issues that arise by comparing memory B cell responses across different antigenic settings^5, 27^. In previous studies, adoptive transfer of clonal BCR transgenic B cells was used to assess antigen-specific B cells in response to LCMV. However, in these models, most if not all transferred cells were eventually depleted^28–30^, thus prohibiting tracking of memory B cell formation over time. To bypass these potential obstacles, we established a LCMV-nucleoprotein (NP)-specific B cell tetramer to identify low-frequency, endogenous, polyclonal antigen-specific B cells at multiple timepoints post-infection (Figure 1A). The NP protein sequence is almost identical between LCMV-WE and LCMV-Docile strains (data not shown), and unlike the glycoprotein does not decline over time in infected cells^31^.

**Figure 1.**
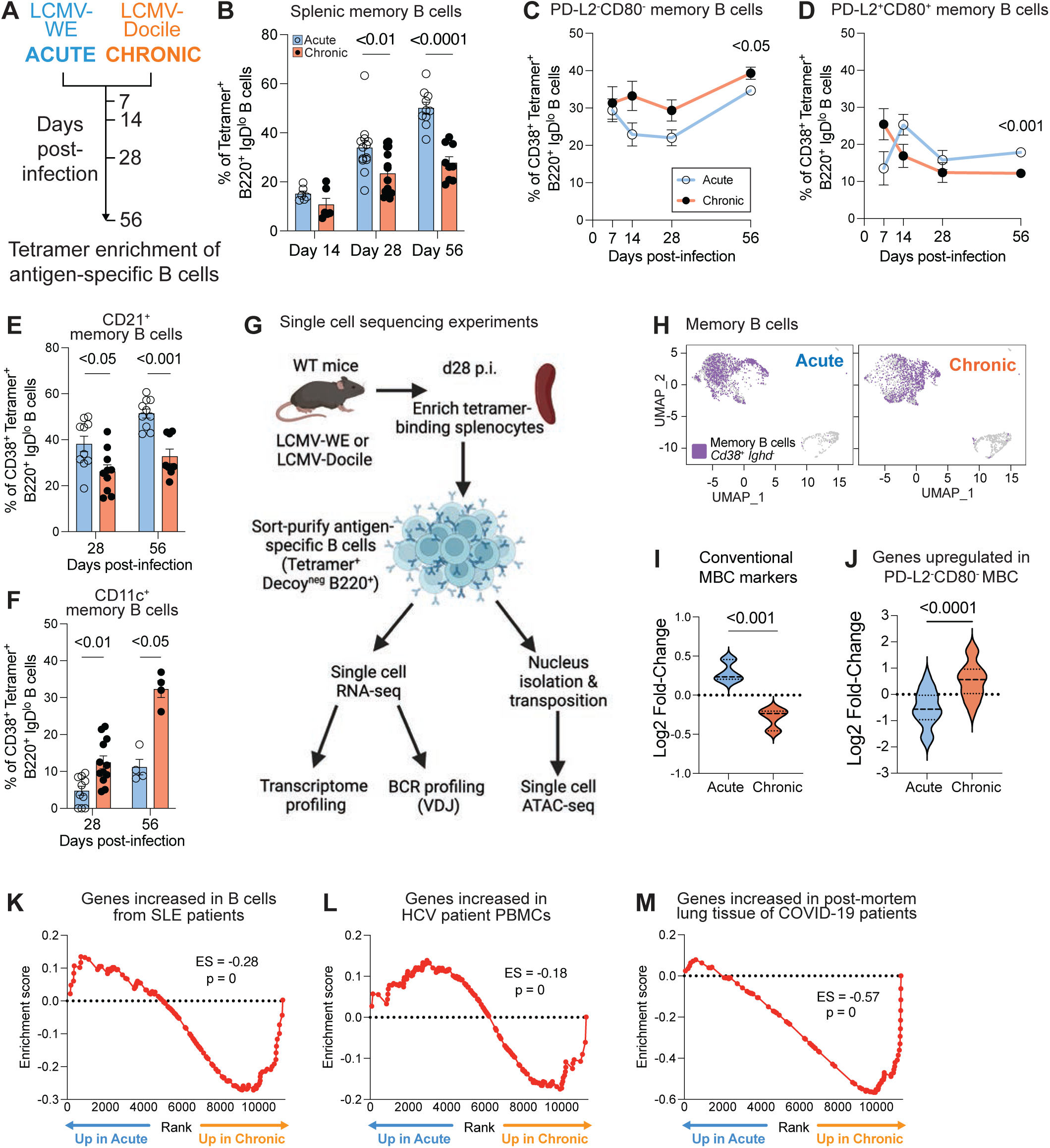
Memory B cells in LCMV-Docile infection adopt phenotypic changes in line with chronic disease contexts in human patients. (A) Schematic of acute (LCMV-WE) or chronic (LCMV-Docile) LCMV infection in C57BL/6 mice. Antigen-specific B cell populations assessed at various timepoints post-infection. (B) Flow cytometric analyses of splenic memory B cells (CD38^+^ % of B220^+^IgD^−^ Tetramer^+^Decoy^−^) in mice at day 14 (WE: n = 6, Docile: n = 6), day 28 (WE: n = 13, Docile: n = 15), or day 56 (WE: n = 10, Docile: n = 9) post-LCMV-WE or LCMV-Docile infection. Combined data from 7 independent experiments. Data represent mean ± SEM. (C and D) Flow cytometric analyses of (C) splenic PD-L2^−^CD80^−^ and (D) PD-L2^+^CD80^+^ within the memory B cell population (B220^+^ IgD^−^ Tetramer^+^ Decoy^−^ CD38^+^) at day 7 (WE: n = 4, Docile: n = 4), day 14 (WE: n = 11, Docile: n = 9), day 28 (WE: n = 9, Docile: n = 8), or day 56 (WE: n = 10, Docile: n = 8) post-infection. Combined data from 7 independent experiments. Data represent mean ± SEM. (E and F) Flow cytometric analyses of (E) splenic CD21^+^ and (F) CD11c^+^ within the memory B cell population at day 28 (WE: n = 10, Docile: n = 10), or day 56 (WE: n = 10, Docile: n = 9) post-LCMV-WE or LCMV-Docile infection. Combined data from 4 independent experiments. Data represents mean ± SEM. (G) Schematic of scRNA-seq and scATAC-seq set-ups. (H) UMAP of scRNA-seq data, with *Cd38*^+^*Ighd*^−^ non-GC cells split by condition shown as purple dots. (I and J) Violin plots representing the log2 fold-change expression of a panel of genes associated with (I) classical memory B cells (*Pdcd1lg2, Cd80, Cr2, Sell, Cxcr5, Ccr7 and Cxcr4*) or (J) PD-L2^−^CD80^−^ memory B cells in memory B cells (*Cd38*^+^*Ighd*^−^ non-GC antigen-specific B cells) isolated from LCMV-WE vs. LCMV-Docile-infected mice. Data represent mean ± SEM. (K, L and M) Gene Set Enrichment Analysis (GSEA) of (K) genes upregulated in B cells from SLE patients compared to healthy donors (GSE30153), (L) genes upregulated in HCV patient PBMCs compared to healthy donors (GSE40184), (M) genes upregulated in SARS-CoV2 positive patient lung tissue (GSE147507) in memory B cells (*Cd38*^+^*Ighd*^−^ non-GC antigen-specific B cells) isolated from B cells generated in LCMV-WE vs. LCMV-Docile infection. Related to Figure S1.

Wild-type C57Bl/6 mice were infected intravenously with either LCMV-WE or LCMV-Docile and antigen-specific memory B cells were investigated over time. Tetramer-binding B220^+^IgD^−^ cells that did not bind the negative control (decoy) tetramer (Figure S1A) were isolated by magnetic enrichment and analysed for quantitative and qualitative changes between acute and chronic infection. Importantly, few antigen-specific B cells were identified from naïve mice (Figure S1A). Splenic memory B cells steadily increased over time in response to acute infection, up to d56 of assessment (Figure 1B). In comparison, in chronic infection there was a significant reduction in frequency of memory B cells observed at d28 and d56 post-infection compared to acute (Figure 1B). As such, the affinity of nucleoprotein-specific memory B cells produced in chronic infection, as assessed by an antigen monomer assay^32^, was decreased compared to those arising from acute infection (Figure S1B). However, GC B cells were significantly increased following chronic compared to acute infection (Figure S1C), as we and others have previously observed^4, 33^, while memory B cell numbers were comparable by d56 (Figure S1C) due to the increased total splenic cellularity in chronic infection^4^. Taken together, it appeared that chronic LCMV-Docile infection induces prolonged GCs (Figure S1C) and hypergammaglobulinemia^4, 34^ but the memory B cell frequency is restrained over time compared to LCMV-WE (Figure 1B).

### Memory B cells in chronic LCMV infection model phenotypic and transcriptomic changes in human chronic infectious disease

Phenotypically defined subsets within the memory B cell population have been linked to functional capacity^35–38^. For example, conventional memory B cells in mice which express the surface markers PD-L2 and CD80 are primed to differentiate into antibody-secreting cells quickly during a recall response^36^. In comparison, “naïve-like” memory B cells that lack expression of both PD-L2 and CD80 have fewer V gene mutations and are more prone to re-populate secondary GCs^36, 39^. In chronic LCMV infection, we found that there was a significant shift between these subsets: the naïve-like subset was increased in chronic LCMV infection (Figure 1C), while the PD-L2^+^CD80^+^ population was decreased (Figure 1D). Similarly, other canonical memory B cell markers were also decreased (Figure S1D). Another layer of memory B cell complexity is observed in human patients across multiple chronic antigen disease contexts, with the emergence of memory B cells that lack CD21 and upregulate CD11c in the peripheral blood^5^. This population was also increased following chronic LCMV infection in vivo, with memory B cells at d28 and d56 significantly downregulating CD21 in both spleen and peripheral blood (Figure 1E Figure S1D-E) and significantly upregulating CD11c (approximately 3-fold increase at d56; Figure 1F), compared to memory B cells induced by acute LCMV infection. Thus, memory B cells formed in response to chronic LCMV infection adopt phenotypic changes commensurate with chronic disease contexts in human patients.

Having established that chronic LCMV infection induced these phenotypic changes, we sought to investigate the extent to which the transcriptome and chromatin landscape was also disrupted by chronic viral infection. We sort-purified antigen-specific B cells at 4 weeks post-infection, a timepoint at which LCMV-WE has been cleared while LCMV-Docile infection persists, and performed multiomic sequencing (Figure 1G). Specifically, we profiled the transcriptome and immunoglobulin diversity via scRNA-seq and assessed chromatin accessibility via scATAC-seq (Figure 1G). Comparison of genes expressed by memory B cells in our scRNA-seq data to those from other published RNA-seq datasets demonstrated that conventional markers of memory^40, 41^ or genes expressed in memory B cells or memory B cell precursors^42, 43^ were decreased, while genes upregulated in PDL2^−^CD80^−^ memory B cells were increased (Figures 1H – 1J, S1F). Thus, memory B cells generated in response to LCMV-Docile infection showed significant changes in gene expression compared to the previously established conventional memory B cell transcriptome. Gene set enrichment analysis (GSEA) of our scRNA-seq dataset demonstrated gene sets from systemic lupus erythematosus (SLE) patient B cells^44^ (Figure 1K), HCV patients^45^ (Figure 1L) and post-mortem lung tissue from COVID-19 patients^46^ (Figure 1M) were enriched in our LCMV-Docile dataset, suggesting that gene signatures in both chronic inflammation and severe viral infection shared parallels with memory B cells in our chronic mice. Collectively, these data also demonstrate the power of this tetramer-based LCMV system to delineate memory B cell fate in vivo in chronic viral infectious disease.

### Distinct epigenetic and transcriptomic signatures define memory B cell subsets

We next used clustering analyses of the scATAC-seq (Figure 2A) and scRNA-seq (Figure 2B) to determine the key transcriptomic or epigenetic changes that ingrain different memory B cell fates during acute versus chronic infection. In particular, this analysis would illuminate whether the heterogeneity seen at the phenotypic level was imprinted by the epigenome. Indeed, 7 unique clusters were identified by scATAC-seq (Figure 2A), and 12 clusters by scRNA-seq (Figure 2B). We first performed cell annotation using Immunological Genome Project (ImmGen) references to determine how these groups related to B cell differentiation states. The smaller, horseshoe-shaped group of scRNA-seq clusters (R3, R9 and R10) were identified as being GC B cells, confirmed by *Fas* transcript expression and the lack of *Cd38* (Figure 2C, S2A). Furthermore, the gene encoding the proliferation marker Ki67 (*Mki67*) was localised to GC clusters, with few cells in the larger group of clusters expressing *Mki67* (Figure S2B). Conversely, memory B cells, defined as antigen-specific B cells expressing *Cd38* and lacking *Ighd*, were localised to the larger set of clusters (Figure S2A). Our dataset excluded the plasma cell population, as cells expressing the genes encoding the plasma cell markers CD138 and Blimp-1, *Sdc1* and *Prdm1* respectively, were scarce (Figure S2C). While our memory B cell definition would preclude analysis of IgD^+^ memory B cells^47^, it was necessary to ensure exclusion of any antigen-specific naïve B cells. To assess fidelity between datasets, we performed a label transfer whereby scRNA-seq annotations were transferred to the scATAC-seq dataset (Figure S2D, S2E). Analysis of merged data revealed that the GC scRNA-seq clusters overlaid with scATAC-seq cluster A5, and in confirmation, A5 was characterized by key loci of genes upregulated by GC B cells, such as *Aicda, S1pr2* and *Fas* (Figures 2D and S2F).

**Figure 2.**
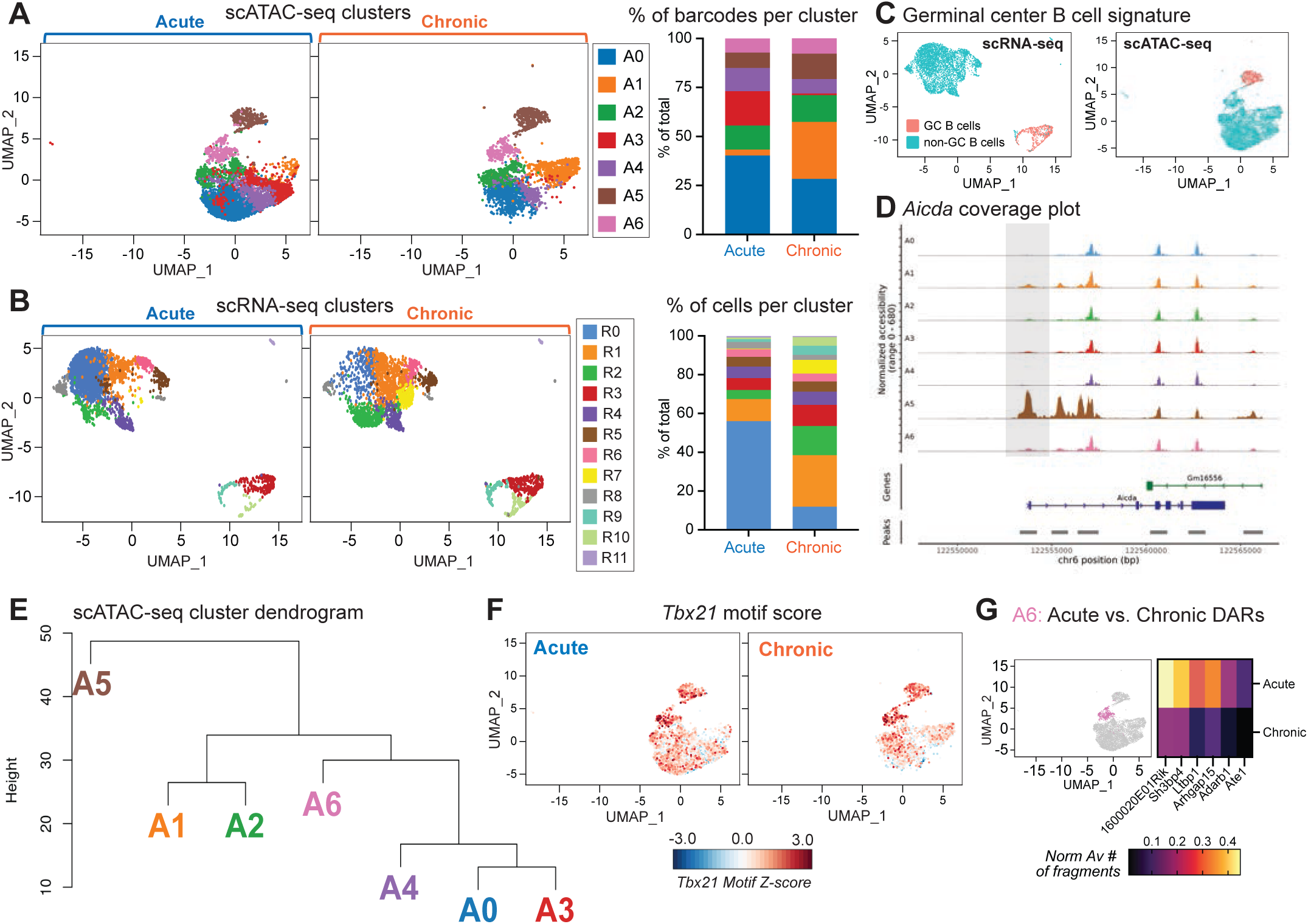
Unique memory B cell subsets emerge following acute versus chronic LCMV infection, defined by distinct transcriptomic and epigenetic profiles. (A) Unsupervised clustering of nuclei from splenic antigen-specific B cells from 2 mice per group visualized using UMAP. Each nuclei is represented by a point and colored by cluster. Graphical representation of percentage of total barcodes per cluster, split by infection type. (B) Unsupervised clustering of splenic antigen-specific B cells from 2 mice per condition visualized using UMAP, split by infection type. Each cell is represented by a point and colored by cluster. Graphical representation of percentage of total cells per cluster, split by infection type. (C) UMAPs of scRNA-seq (left) and scATAC-seq (right) data, with GC B cells (pink) and non-GC B cells (aqua). (D) Coverage plot showing chromatin accessibility at the *Aicda* gene region, split by cluster. (E) Dendrogram showing the relationships in chromatin accessibility between the scATAC-seq clusters. (F) UMAP of *Tbx21* motif enrichment, with Z-score representing the sums of cut sites per cell which fall within all the peaks associated with the *Tbx21* motif, split by condition. (G) UMAP plot with cluster A6 highlighted. Heatmap of the DARs between acute and chronic condition in cluster A6. Related to Figure S2, Figure S3 and Figure S4.

We next assessed the shift in clusters in chronic, compared to acute, LCMV infection. All clusters were detectable in both acute and chronic conditions (Figures 2A and 2B), and the location of chromatin peaks was distributed equivalently along the genome independently of infection condition or cluster (Figure S2G). However, there were clear changes in the distribution of clusters within each infectious condition. In the scATAC-seq dataset, cluster A1 was greatly expanded in chronic infection, whereas cluster A3 had become barely detectable (Figure 2A). Given that one cluster expanded in chronic infection whilst another diminished, we asked whether these clusters were closely related and may in fact just be reflective of a small shift in the chromatin landscape between infections. We performed a pseudo-bulk analysis of each cluster and calculated a distance matrix to visualize the hierarchical relationship between clusters. Despite the paired change in these clusters, the chronic (A1) and acute (A3) expanded clusters were highly distinct from one another (Figure 2E).

To understand whether the observed loss of a memory B cell subset correlated to the decrease in conventional memory B cell phenotype observed in Figure 1, we interrogated our scRNA-seq dataset. Cluster R0 dominated the acute memory B cell population, constituting approximately half of all tetramer-specific cells in the acute-infected mice, whilst R7 was barely detectable. However, this distribution was notably reversed in chronic infection, with cluster R0 substantially diminished (Figure 2B). Cluster R0 had increased expression of *Cd55*, which serves to protect host cells from complement-mediated damage, *Fcer2a* which encodes CD23, and the transcription factor *Klf2*, which regulates cell trafficking via upregulating CD62L^48^, the gene for which (*Sell*) was also upregulated in R0 (Figure S3A). Several transcription factors previously associated with memory B cell formation were also decreased in chronic infection (Figure S3B-C). Given the shift away from a conventional memory B cell identity following chronic LCMV infection, we assessed isotype distribution and somatic hypermutation (SHM) frequency across clusters by scVDJ-seq (Figures S3D and S3E). While there was little change in isotype distribution in antigen-specific memory B cells in acute and chronic infection (Figure S3D), there was a large reduction in unmutated R0 cells, paired with an increase in somatically mutated GC B cells (Figure S3E). Taken together, clustering analysis confirmed that chronic viral infection imparted discernible, sustained changes to the memory B cells, with the emergence of distinct memory B cell subsets and the diminution of memory B cells with a conventional transcriptomic identity.

### T-bet-expressing memory B cells are not rewired by chronic LCMV infection

One of the defining features of memory B cell changes during chronic infectious disease has been the increased prominence of a subset expressing T-bet, CD11c (encoded by the gene *Itgax*) and CXCR3 in human peripheral blood^5^. However, T-bet-expressing memory B cells are also induced in acute infection^13, 14, 17, 18, 49^. One key question that has remained unanswered is whether the T-bet-associated memory B cell subset which arises in chronic disease is distinct from T-bet-expressing memory B cells produced in acute immune responses. To answer this question, we first asked whether this population could be uniquely resolved as a distinct population in our multiomic dataset. ATAC cluster A6 showed enrichment of *Tbx21* motif (Figure 2F) and had increased chromatin accessibility within genes associated with T-bet-expressing memory B cells, including *Tbx21*, *Itgax* and *Zeb2* (Figures S4A – S4C). Notably, A6 overlapped with cluster R5 in the merged dataset (Figure S2E), which correspondingly showed increased expressed of T-bet-associated genes (*Tbx21*, *Cxcr3*, *Itgax*, *Cd80*, *Zeb2*), whereas other memory markers such as *Cr2, Hhex and Pdcd1lg2* had a wider distribution across clusters (Figures S4D, and S4E). The relationship between A5 and A6 (Figures 2E, S4A), as well as the SHM frequency (Figure S3D), suggested that T-bet-expressing cells at week 4 post-LCMV infection may be derived from both GC-dependent in addition to GC-independent pathways^17, 50^. In summary, both acute and chronic LCMV infection were able to induce a *Tbx21-*expressing memory B cell subset, with shared transcriptomic profiles.

To determine whether chronic infection leaves ‘epigenetic scars’^51^ on this population, we compared the epigenetic signature between acute versus chronic barcodes in cluster A6 (Figure 2G). Only 6 differentially accessible regions (DARs) were detectable between conditions, indicating that *Tbx21*-expressing subset had a similar epigenetic profile regardless of acute or chronic infectious context. Furthermore, comparison of the transcriptome in acute versus chronic infection identified only two significantly different genes, *Ly6c* and *Ifi2712a*, which also showed increased expression across other clusters. Thus, chronic LCMV infection did not impart unique changes to the T-bet^+^ population that may alter the characteristics of this memory B cell subset in chronic infection.

### Chronic LCMV infection induces an ISG-enriched memory B cell population

We then turned our attention to the unresolved question: what are the defining molecular features of memory B cells that are uniquely expanded during chronic viral infection? To investigate how chronic infection remodels the memory B cell population, we examined the attributes of the scATAC-seq and scRNA-seq clusters that were expanded in chronic, compared to acute, viral infection. There was a clear expansion of cluster A1 in chronic infection (Figure 2A). To identify unique properties of the chronic-emergent A1 cluster, we compared its chromatin landscape to all other non-GC clusters. This analysis identified 39 distinct chromatin regions that were open in A1 (Figures 3A and 3B). These regions were enriched for binding motifs of transcription factors important for B cell biology, as assessed by HOMER^52^ (which tests for motif enrichment in differentially accessible regions; Figure 3C) or chromVAR^53^ (which tests for significant differential motif activity between clusters; Figure S5A). For example, several with known roles in regulating B cell transcriptional programs (e.g., E2A, PU.1, Foxo1) and negatively regulating GC B cells (Bhlhe40; corresponding increased transcriptional expression in chronic memory B cells shown in Figure S5B)^54^ were enriched in areas of increased chromatin accessibility in chronic memory B cells.

**Figure 3.**
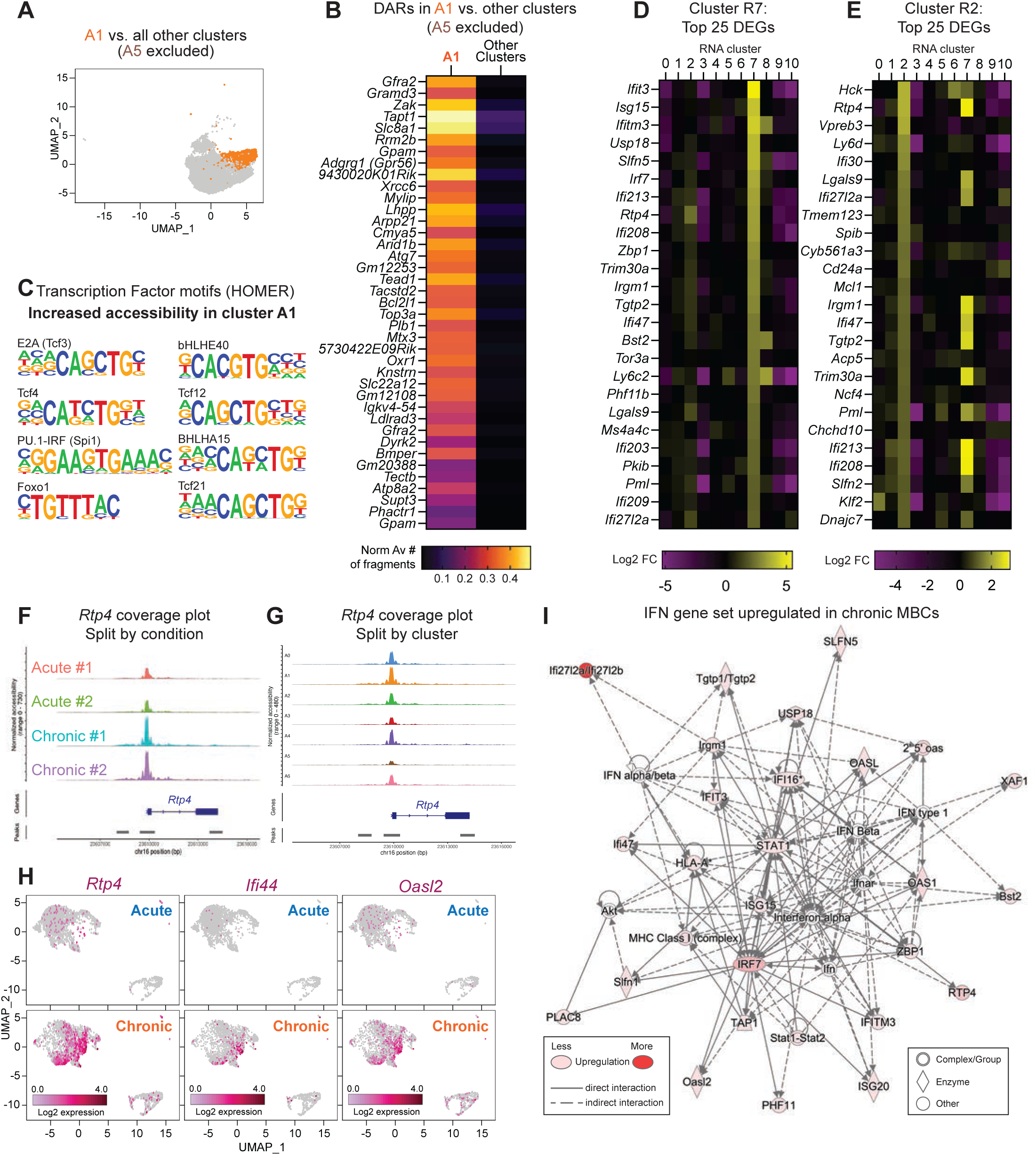
Identification of distinct chronic infection-expanded Rtp4 and ISG-associated subsets. (A) UMAP plot with cluster A1 highlighted (excluding cluster A5). (B) Heatmap of the DARs in cluster A1 compared to all other clusters (excluding cluster A5). (C) Representative selection of transcription factor binding motifs enriched in cluster A1 vs. all other clusters (excluding A5), generated using HOMER. (D and E) Heatmaps of the top 25 DEGs in (D) cluster R7 or (E) cluster R2 (excluding Ig genes), compared to all other clusters. (F and G) Coverage plots showing chromatin accessibility at the *Rtp4* gene region, split by (F) condition, and (G) split by cluster. (H) UMAPs of gene expression of *Rtp4*, *Ifi44* and *Oasl2* split by condition, with positive expression represented by pink dots. (I) IPA analysis of differentially expressed genes within the IFN pathway upregulated in memory B cells (*Cd38*^+^*Ighd*^−^ non-GC) from LCMV-Docile-infected mice. Related to Figure S5.

Next, we interrogated the chronic-emergent clusters from the scRNA-seq dataset, clusters R7 and R2 (Figures 3D and 3E). Cluster R7 was almost entirely unique to mice responding to chronic infection, with only ∼0.2% of antigen-specific B cells in acute infection populating this cluster. 20 out of the top 25 upregulated differentially expressed genes (DEGs) in cluster R7 were ISGs: *Ifit3, Isg15, Ifitm3, Isg20, Usp18, Slfn5, Irf7, Ifi213, Ifi208, Trom30a, Irgm1, Ifi47, Bst2, Tor3a, Ly6c2, Phf11b, Lgals9, Ifi203, Ifi209, Ifi27l2a* (Figure 3D). Cluster R2 shared many of the top DEGs from R7, including *Lgals9, Ifi27l2a, Ifi47, Ifi203, Ifi208* and *Rtp4* (Figure 3E). Both of these scRNA-seq subsets overlapped with A1 from the scATAC-seq data, although Cluster R7 had a broader distribution (Figure S2E). The high expression of *Rtp4* in both R2 and R7 was particularly of note: Rtp4 is IFN-I-inducible and is a regulator of type I IFN responses^55^. Chromatin accessibility was significantly increased at the *Rtp4* promoter region in antigen-specific B cells in chronic compared to acute infection (Figure 3F), particularly in the chronic-expanded cluster A1 (Figure 3G). The increased accessibility and increased expression of *Rtp4* also appeared to be functionally consequential, with the majority of Rtp4-regulated genes upregulated in memory B cells during chronic infection^55^ (Figure 3H and S5C). Additionally, Ingenuity Pathway Analysis (Figure 3I) confirmed that chronic memory B cells were enriched for an IFN gene set. This ISG signature appeared to be concentrated in clusters that were expanded in chronic infection (*Ifi44 and Oasl2,* Figure 3H; *Irf7,* Figure S5D), although some genes did have an overall general increase in expression (e.g. the IFN-inducible *Ifi2712a*, Figure S5D). Thus, the scRNA-seq and scATAC-seq data identified a transcriptomically and epigenetically unique type of memory B cell expanded in chronic LCMV infection, which was dominated by an ISG signature and was independent of the previously described *Tbx21*-expressing memory B cell subset.

### Type I IFN is a key driver of phenotypic and epigenetic changes to memory B cells during chronic viral infection

The timing and magnitude of IFN is an important determinant for whether viral infections will progress to severe disease^56^. A rapid but transient peak of IFN-I in the first day of infection enables the cellular response to effectively clear the virus. In contrast, low and/or delayed IFN-I induction leads to persisting viral loads, an ineffective and eventual dysregulated immune response, and increased morbidity^6, 56–58^. Given the ISG signature in the unique chronic-emergent subsets at 4 weeks post-infection in both the scRNA-seq and scATAC-seq analysis (Figure 3), we hypothesized that IFN may be instructing identity and/or driving maintenance of the chronic memory B cell subset. In line with this hypothesis, IFN*α* response and IFN*γ*response hallmark gene sets were enriched in the memory B cells in chronic LCMV infection (Figure 4A). Thus, we assessed whether either type I or type II IFN was a key driver of chronic memory B cell identity. Wild-type mice infected with LCMV were administered a blocking antibody to either IFNAR-1 or IFN*γ* every 3 days from d2 to d14 post-infection (Figure 4B). IFNAR-1-blocking induced qualitative and quantitative changes to the memory B cell population, while IFN*γ*-blocking did not (Figures 4C – 4E). While the frequency of antigen-specific IgD^−^ B cells was similar with or without IFN blocking treatment (Figure S6A), the frequency of memory B cells, and CD11c^+^, PD-L2^+^CD80^+^ and PD-L2^−^CD80^+^ memory B cell subsets were increased, while CD21 expression remained the same (Figures 4C – 4E, Figures S6B – S6D). Overall, activated B cell numbers were also increased (Figure S6E), which may be due to inhibition of IFN-mediated B cell-depleting mechanisms described in previous studies using adoptive transfer models^28–30^.

**Figure 4.**
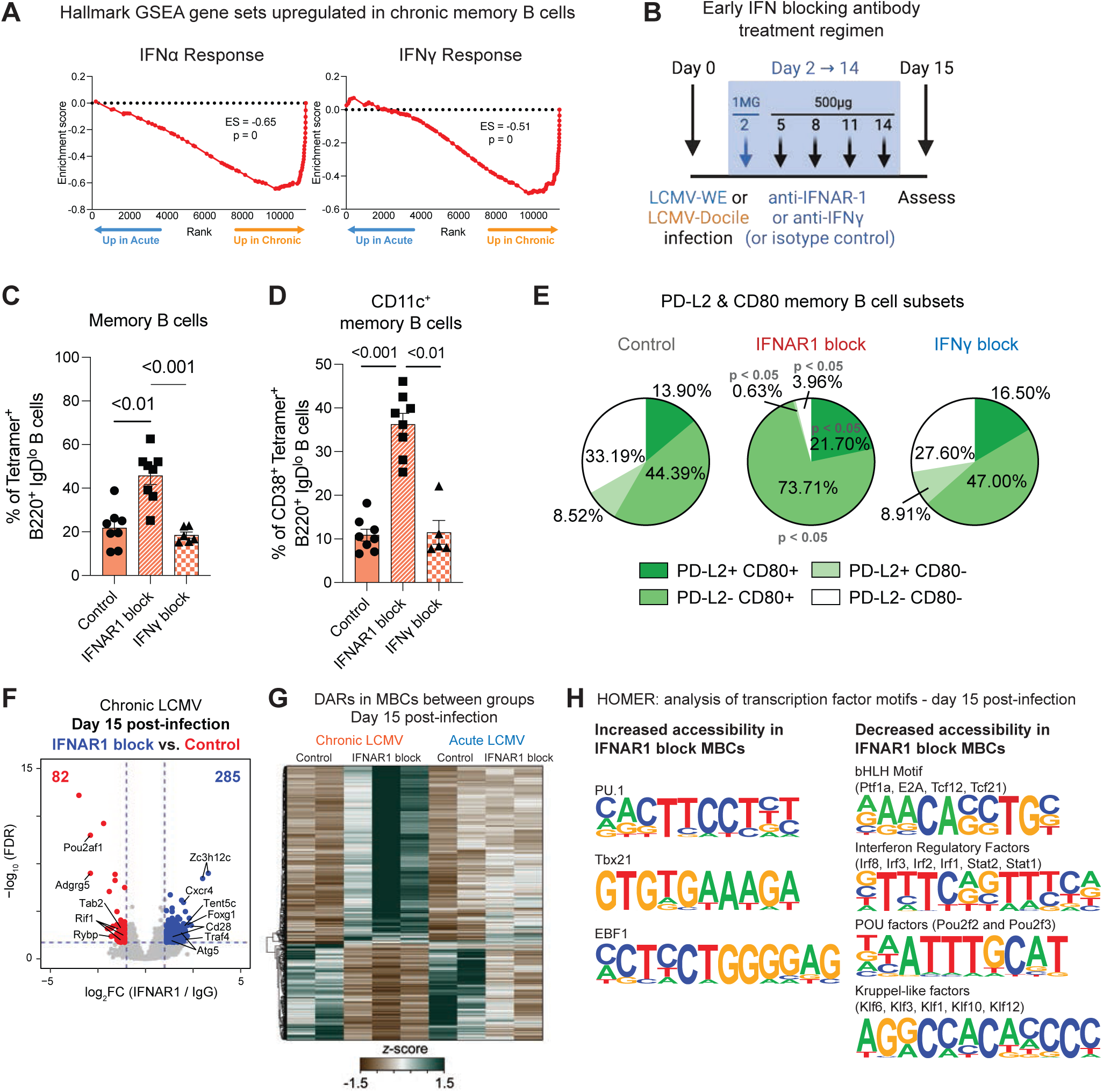
Type I IFN is a key driver of phenotypic and epigenetic changes to memory B cells during chronic viral infection. (A) GSEA of hallmark IFNα response (left) and IFNγ response (right) gene sets in memory B cells (*Cd38*^+^*Ighd*^−^ non-GC) isolated from antigen-specific B cells generated in LCMV-WE vs. LCMV-Docile infection. Experimental design shown in Figure 1. (B) Schematic of early type I or II IFN blocking treatment cohort and timepoint. (C, D and E) Flow cytometric analyses at day 15 post-infection of (C) antigen-specific memory B cells (B220^+^IgD^−^Tetramer^+^Decoy^−^CD38^+^), (D) CD11c^+^ frequency within memory B cells, and (E) PD-L2 and CD80 memory B cell subsets in mice treated with anti-IFNAR1 (MAR1-5A3) (n = 8), anti-IFNγ (XMG1.2) (n = 5) or IgG control (n = 8) at day 15 post-infection. Combined data from 2 independent experiments. Data represents mean ± SEM. (F) Volcano plot of DARs in memory B cells in IgG control (red) vs. anti-IFNAR1 blocking treatment (blue) in LCMV-Docile infection at day 15 post-infection. (G) Heatmap showing DARs in memory B cells in LCMV-WE and LCMV-Docile infection, with IgG control, anti-IFNAR1 or anti-IFNγ blocking treatment at day 15 post-infection. (H) Representative selection of transcription factor binding motifs with increased (left) or decreased (right) chromatin accessibility in memory B cells following early IFNAR1-blocking treatment generated using HOMER. Related to Figure S6.

We next tested whether IFN-I blockade could induce extensive remodelling of the chromatin landscape within memory B cells. Antigen-specific CD38^+^IgD^−^ memory B cells were sort-purified at day 15 post-infection after blocking antibody treatment, and ATAC-seq was performed. The chromatin landscape of memory B cells was largely remodelled in chronic mice treated with IFNAR-1 blocking antibody, compared to mice treated with control antibody, with 616 DARs identified (Figure 4F and 4G). In stark contrast, IFNAR-1 blockade during acute LCMV infection induced far fewer changes in the memory B cell chromatin landscape (Figure 4G). In chronic mice, gains in accessibility in memory B cells were associated with access to transcription factor motifs such as *Pu.1*, *Ebf1* and *Tbx21* (Figure 4H), the latter likely responsible for the increased in CD11c expression (Figure 4D). While there were fewer changes associated with decreased accessibility, the transcription factor binding motifs in these decreased DARs were concordant with those that had been identified in the ISG chronic memory B cell subset (Figures 3C and S5A). In line with IFN response regulation, these included IFN regulatory factors (Figure 4H); in addition, Tcf3, Tcf12, Tcf21, Ptf1a and POU factors (Figures 4H, 3C and S5A). Kruppel-like factors, which can play a role in increasing accessibility to cofactors^59^, were also detected in these DARs. Together, these data suggest that IFN-I signaling is a critical factor governing memory B cell identity at the epigenetic and phenotypic level in chronic LCMV infection.

### Viral load is not a sole driver of chronic memory B cell phenotype

The ongoing persistence of virus and viral load has been linked with changes to memory B cell phenotype and incomplete immune protection in HIV patients^19, 60^. While previous studies have shown that although the majority of memory B cells are formed within the first week, they can also continually form throughout an ongoing response^34, 43, 47, 61–63^. We therefore expected that the ongoing presence of virus was an important determinant in promoting chronic memory B cell attributes, in addition to the role of the early microenvironment factors observed in Figure 4. Yet, patients on anti-retroviral therapy do not completely restore immune memory populations^64–67^, suggesting that there may be a time-based limitation on the establishment of long-lived memory B cell identity during chronic viral infection. We therefore tested the timing of memory B cell formation and the extent to which viral load shaped memory B cell development and maintenance in chronic infection. We first investigated the in vivo kinetics of memory B cell formation and longevity in acute versus chronic viral infection with 5-bromo-2’-deoxyuridine (BrdU) labelling experiments. A defining characteristic of long-lived memory B cells is that they become quiescent^7^ and thus should retain BrdU if formed during the labelling period. Infected mice were administered BrdU over 3-day windows (Figure 5A). BrdU uptake and long-term retention in specific cell populations was examined across the course of infection (Figure 5B-D). Our data indicated that the majority of BrdU^+^ memory B cells detected at d56 post-infection were generated early during the course of infection (BrdU-labelled D5-7; Figure 5B). LCMV-WE-induced memory B cells generated in this early time frame were stable over time, while BrdU^+^ chronic memory B cells progressively declined (Figure 5B). This was likely due to continual viral presence-induced proliferation, as there was an increase of BrdU^+^ memory B cells in chronic infection when BrdU was administered in the days immediately prior to d56 (data not shown).

**Figure 5.**
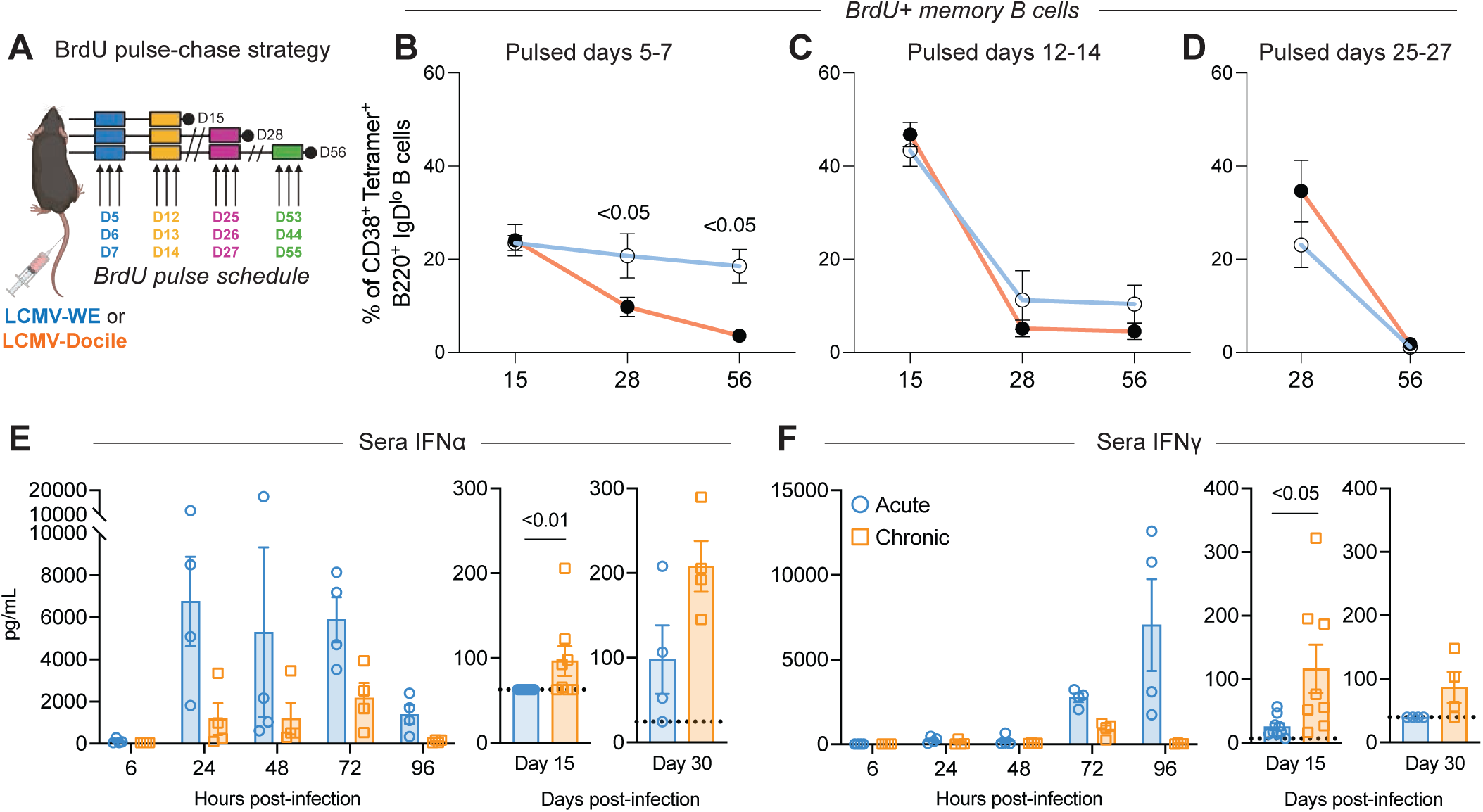
Chronic LCMV infection had delayed induction of type I and type II IFN, but sustained IFN exposure and continued proliferation of memory B cells late into the course of infection. (A) Schematic of BrdU treatment groups and timepoints following LCMV-WE or LCMV-Docile infection. (B, C and D) Flow cytometric analyses of BrdU^+^ frequency within memory B cells (B220^+^IgD^−^ Tetramer^+^Decoy^−^CD38^+^) at various timepoints post-infection in mice infected with LCMV-WE or LCMV-Docile and treated with BrdU at (B) days 5-7 post-infection, (C) days 12-14 post-infection or (D) days 25-27 post-infection. Combined data from 3 independent experiments. Data represents mean ± SEM. (E and F) LegendPlex assay results of sera (E) IFNα or (F) IFNγ at various timepoints post-infection with LCMV-WE (n = 4-8) or LCMV-Docile (n = 4-8). Combined data from 3 independent experiments. Data represents mean ± SEM. Related to Figure S6.

In line with the essential role of IFN-I signaling for viral control early in the course of infection^68–70^, we found that IFNAR-1 blockade increased viremia in chronic LCMV infection at d15 post-infection (Figure S6F). We therefore asked whether elevated viremia, upstream of IFN production, may drive changes to memory B cell identity during chronic LCMV infection, rather than IFN itself. To assess this possibility, we used the antiviral pyrazinecarboxamide-derivative Favipiravir, which is broadly active against a wide variety of RNA viruses including LCMV^71^. We administered Favipiravir daily from d2-8 post-infection and culled mice at d12 to investigate whether memory B cell subsets were reliant on the presence of elevated viremia (Figure S6G). However, although Favipiravir treatment significantly lowered viral titers (Figure S6H), the distribution of memory B cell subsets did not fundamentally change (Figure S6I-S6K). This was true also when treatment was performed at 4 weeks post-infection (data not shown). Thus, the chronic memory B cell phenotype was not solely driven by sustained high viremia.

### There is a critical window in which IFN-I exposure directs establishment of the memory B cell chromatin landscape during chronic viral infection

Our results thus far showed that the memory B cell phenotype and chromatin landscape could be shaped by IFN during the first two weeks of infection. However, it was somewhat surprising that IFN blocking administration had a limited effect on the chromatin landscape of memory B cells during acute infection. Furthermore, we hypothesized that IFN may be elevated in chronic infection later in the response, given the ISG signature in memory B cells was prominent at d28 post-infection. We therefore compared the kinetics of IFN production in LCMV-WE vs. LCMV-Docile infection. While previous studies have compared IFN induction in LCMV-Armstrong versus LCMV-Clone13 in fine detail^56, 57, 72^, direct comparison of WE and Docile beyond the first two days post-infection^68^ was lacking. As previously described, LCMV-WE induced a high wave of IFN*α*by 24hr post-infection, which is critical to mount an efficient immune response^6^. In comparison, IFN*α* in mice infected with LCMV-Docile was delayed and did not reach the same peak as LCMV-WE before declining by d4 post-infection (Figure 5E). Similar differences between WE and Docile strains were observed with IFN*γ*kinetics (Figure 5F). While IFN induction was delayed in chronic infection, the amount of IFN in LCMV-Docile mice at d15 and d30 post-infection was significantly increased over LCMV-WE and at a similar concentration to that observed at d4, suggesting a low level of sustained IFN in chronic infected mice (Figure 5E & 5F).

Given the sustained presence of IFN in chronic infection, we asked whether the continual low exposure of IFN-I was critical for the maintenance of the chronic memory B cell identity. To investigate, we treated mice with IFN-I blocking antibody late in the course of an established infection, from d28 to d40 post-infection (Figure 6A). We also administered an IFN*γ* blocking antibody given that there was also a sustained increase in IFN*γ*over time (Figure 5F), and the possibility that IFN*γ*could be inducing the ISG signature detected in our chronic memory B cell subset as has been shown in CD4^+^ T cells^73^ (Figure 6A). However, the use of these blocking antibodies did not significantly alter memory B cell frequency or phenotype (Figures 6B – 6E). Therefore, memory B cell phenotype was impervious to phenotypic change upon IFN blocking antibody administration late in the primary response to chronic LCMV infection. In agreement with the phenotypic data, we identified only 14 DARs between mice that received the control treatment and the IFNAR-1 blockade at day 41 post-infection (Figure 6F – 6I, Figure S6L). Therefore, there is a critical window early in infection in which IFN-I can govern memory B cell identity. Collectively, these experiments provide critical insights into how changes in IFN dynamics during early viral infection can redirect memory B cell development in addition to IFN-mediated effects on viral persistence and severity of disease.

**Figure 6.**
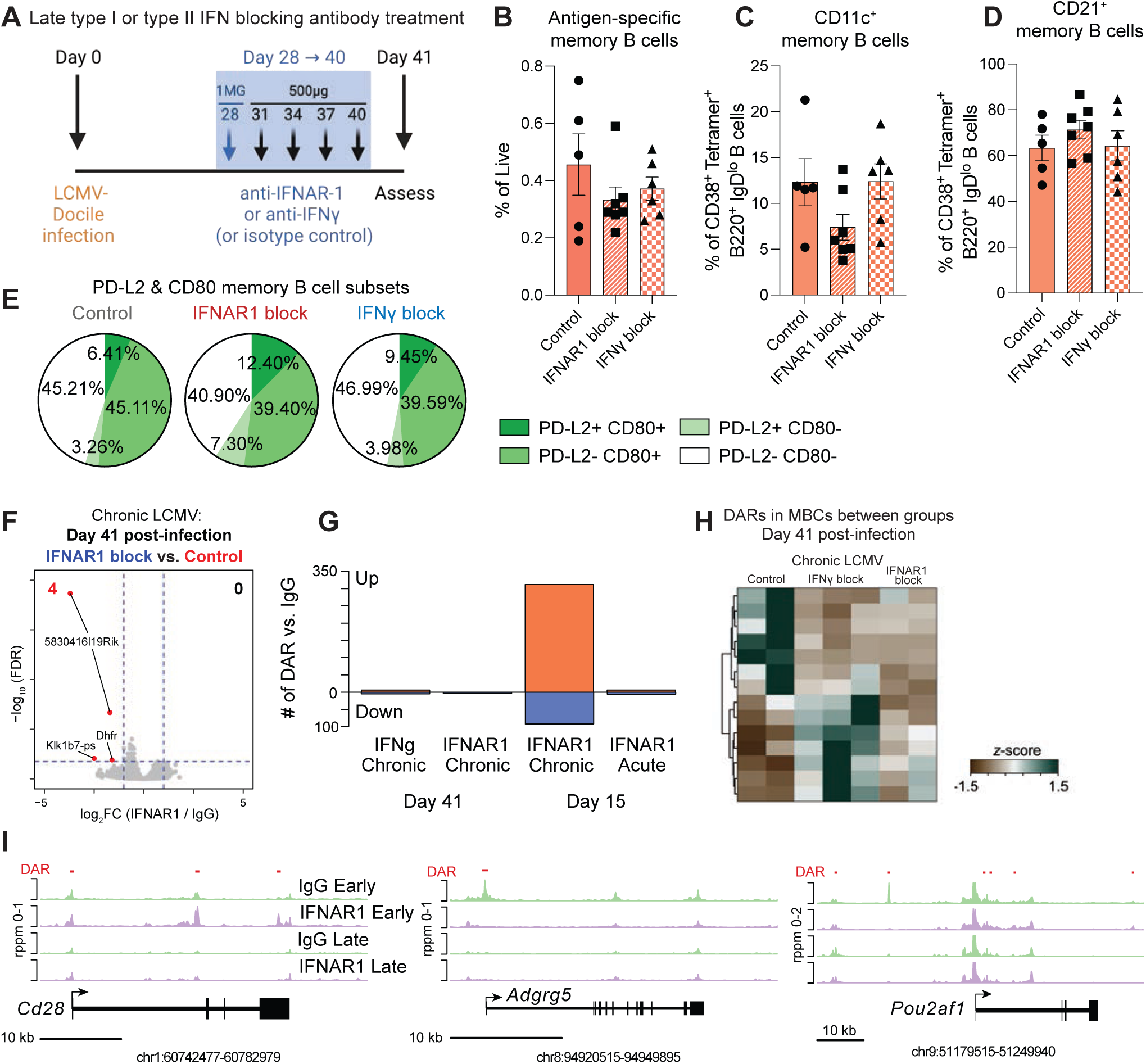
Phenotype and epigenetic profile of memory B cells were impervious to change upon IFN blocking antibody administration late in the primary response to LCMV-Docile infection. (A) Schematic of late type I or II IFN blocking treatment cohort and timepoint. (B, C and D) Flow cytometric analyses of (B) antigen-specific memory B cells (B220^+^IgD^−^ Tetramer^+^Decoy^−^CD38^+^), (C) CD11c^+^ frequency within memory B cells, and (D) CD21^+^ frequency within memory B cells in mice treated with anti-IFNAR1 (MAR1-5A3; n = 7), anti-IFNγ (XMG1.2; n = 6) or IgG control (n = 5) at day 41 post-infection. Combined data from 2 independent experiments. Data represents mean ± SEM. (E) Flow cytometric analyses of PD-L2 and CD80 memory B cell subsets in mice treated with anti-IFNAR1 (MAR1-5A3; n = 3), anti-IFNγ (XMG1.2; n = 3) or IgG control (n = 2) at day 41 post-infection. Combined data from 2 independent experiments. Data represents mean ± SEM. (F) Volcano plot of DARs in memory B cells in IgG control (red) vs. anti-IFNAR1 blocking treatment (blue) in LCMV-Docile infection at day 41 post-infection. (G) Graph representing number of DARs in memory B cells in treatment groups versus IgG control at day 15 and day 41 post-infection. (H) Heatmap showing DARs in memory B cells in IgG control, anti-IFNAR1 or anti-IFNγ blocking treatment in LCMV-Docile infection at day 41 post-infection. (I) Coverage plots showing chromatin accessibility at the *Cd28* (left), *Adgrg5* (center) and *Pou2af1* (right) gene regions, split by treatment group. Related to Figure S6.

## DISCUSSION

Here, we set out key findings which define how the chronic memory B cell identity diverges from memory B cells responding to acute viral infection. Integration of scATAC-seq and scRNA-seq revealed that the diversity in the memory B cell population is wired by the epigenome. While the majority of clusters were present in both acute and chronic LCMV infection, a clear expansion of a cluster characterised by ISGs was identified in chronic infection that was both independent of the *Tbx21*-expressing cluster and seemingly at the expense of a canonical memory B cell subset. The chromatin landscape of chronic memory B cells was established early in the primary response, with IFNAR-1 signaling a key regulator of memory B cell subset identity. After this period, modulation of viral load or IFN blockade had little effect on chromatin accessibility or phenotypic distribution of subsets, indicating that memory B cells were epigenetically stable. Thus, plasticity was not inherent within the established memory B cell pool, even with continual proliferation induced by persisting virus, limiting the potential to therapeutically convert memory B cell subsets. Collectively, our research illuminates that the early microenvironment not only charts a course for the severity of the disease, but also determines the trajectory of chronic memory B cell subset development.

Our data therefore suggests a limited window of opportunity to enact large-scale changes to the memory B cell chromatin landscape. This is congruent with previous studies demonstrating that the bulk of memory B cells forms early after immunization of mice with a model antigen^61^. However, given that fluctuations in viral load have been associated with similar fluctuations of atypical memory B cells in HIV patients, we expected that the ongoing presence of virus would enable remodelling of the memory B cell population by administration of an antiviral or cytokine blocking experiments. Yet, despite the ongoing viral load and ongoing memory B cell proliferation, our data showed that memory B cells were epigenetically stable after the first month of chronic viral infection. This may in fact explain why HIV patients on anti-retroviral therapies are less likely to completely recover their ability to generate effective immune protection, even if memory B cell phenotype has been partly restored to resemble healthy patients^20, 60, 64–67, 74^. Thus, memory B cell subset identity may be imprinted in a ‘hit and run’ manner, in which the distinct epigenetic signatures of each cluster are established in the initial phase of the response.

As such, chronic infection did not largely change the core epigenetic identity of many of the memory B cell subsets formed, but instead expanded or contracted their overall representation. However, by avoiding a priori assumptions of which phenotypic markers chronic memory B cells should possess, our data revealed that there was a clear expansion of a unique subset enriched with an ISG signature in chronic infection. Similar enrichment of ISGs has been identified in analyses of B cell populations in HIV patients^75, 76^. Notably, this chronic-emergent subset was independent of the *Tbx21-*associated, CD11c-expressing subset that dominates descriptions of ‘atypical’ or ‘age-associated’ B cells in humans and mice^16, 77–79^. In fact, it is possible that the ISG signature is highly expressed in the chronic memory B cell subset in part due to the absence of T-bet expression^73^. In contrast to the chronic-emergent subset, we corroborate that Tbet^+^CD11c^+^ cells are not uniquely reliant on a chronic viral infection to be formed, nor does chronic infection impart major changes on their chromatin landscape. In addition to previously established positive regulators IFN*γ*, TLR7 and IL-21^9, 16, 80–82^; we also show that formation of this subset can be promoted by IFNAR-1 blockade. The function of this subset appears to be context-dependent, ranging from protective^83, 84^ to unnecessary for long-lived immunity^16^ to detrimental in human patients responding to different types of disease^85^. Thus, depending on whether this subset is functional (e.g. in malaria) or dysfunctional (e.g. in HIV or systemic lupus erythematosus), our data suggests a therapeutic strategy for modulation.

The kinetics and magnitude of IFN-I responses during viral infection is known to have an important role in disease pathogenesis across various contexts of viral infection^69, 86, 87^. Accumulating evidence points to delayed induction of a low amount of IFN-I being a key determinant in the severity and persistence of viral infection. This has particularly come to the fore during COVID-19^6, 58^ and the postulated mechanistic relationship between IFN and severity of disease in response to SARS-CoV-2 infection and Long COVID^8, 88, 89^. Our study gives critical insight into how memory B cells may be altered by this difference between acute (high, quick load of IFN-I) versus chronic (delayed, low but sustained IFN-I levels). The IFN-I-induced shaping of the memory B cell compartment may explain why patients with autoantibodies to IFN can form immune memory to vaccination but do not generate adequate memory protection during SARS-CoV-2 infection^90, 91^. It also suggests that IFN-I therapy^69, 89^ may have a significant impact on memory B cell development. It is possible that these differences also lead to the induction of CD21^−^ CD27^−^ ‘double negative’ cells in Long COVID^92^, thus acting as a biomarker of the early microenvironment during the nascent immune response. Thus, studying the chromatin landscape at single-cell resolution in vivo can enable the identification of new biomarkers of the critical early immunological events that govern the effectiveness of the host response and thus disease severity.

Beyond vaccines, few strategies exist to exert influence over the development of memory B cells. This is a significant gap in our capabilities not only for treating non-vaccine-preventable infections, but also for immunodeficiencies in which patients cannot form adequate humoral memory. Our research demonstrates that antigen-specific B cell subset distribution during infection is underpinned by distinct epigenetic states, and that it is possible to remodel memory B cells to promote or repress certain subsets. In addition, we have previously shown that the deletion of an epigenetic regulator can alter memory B cell subset distribution and hence their function during a recall response^93^. Given that epigenetic factors can be targeted in vivo, such as with small molecule inhibitors^4^, these findings portend the development of new ways to therapeutically target memory B cells. Development of this strategy and identification of druggable targets will be further enhanced by illuminating the epigenome – that is, the repressive or activation marks that regulate chromatin landscapes – that may enable longevity and rapid responsiveness of memory B cells in productive immune responses or promote dysfunction during disease.

In summary, single-cell resolution has illuminated the shared and divergent splenic memory B cell populations during chronic versus acute viral infection. Furthermore, our research shows how chromatin landscapes underpinning memory B cell subsets can be reshaped through targeting IFN-I, thus initiating a new therapeutic scope for promoting sought-after efficacious memory B cell subsets or repressing the formation of dysfunctional cells in disease.

## ACKNOWLEDGEMENTS

We thank David Tarlinton and Liam Kealy for critical reading of this manuscript; members of the Good-Jacobson lab for technical assistance; and staff of the Monash University FlowCore, Animal Research Platform, Micromon, Monash Health Translational Precinct, David Powell and the Monash University Bioinformatics Platform.

## AUTHOR CONTRIBUTIONS

KLG-J conceived the study; KLG-J, LC and IAP designed research; LC, JP, LM, AN, EESP and HX performed research; KLG-J, LC, KJLJ, EESP, MG and CDS analyzed data; IAP, MOK and NLG provided intellectual input; JJT, LM, CS and SG provided reagents and technical expertise; and KLG-J and LC wrote the manuscript.

## METHODS

### Resource availability

#### Lead contact

Further information and requests for resources and reagents should be directed to the lead contact Kim Good-Jacobson.

### Materials availability

Please contact Stephanie Gras or Justin Taylor for biotinylated proteins generated in this study.

### Experimental model and subject details

#### Mice and viral infections

C57Bl/6 mice were maintained at the Monash Animal Research Platform. Both males and females, over 6 weeks old, were used in this study. The Monash Animal Ethics Committee approved all procedures. Mice were culled at indicated time intervals.

*Acute LCMV infection:* intravenous (i.v.) injection of 3 x 10^3^ PFU of LCMV-WE.

*Chronic LCMV infection:* i.v. injection of 2 x 10^6^ PFU of LCMV-Docile.

#### BrdU administration

5-bromo-2’-deoxyuridine (BrdU; Sigma-Aldrich) was diluted to 5mg/ml in sterile PBS. 200uL (1mg) was injected intraperitoneally into each recipient mouse at indicated labelling time points in order to label proliferating cells. PBS alone was used for unlabelled control mice.

#### Favipiravir treatment

Favipiravir (Biorbyt) was diluted in sterile PBS 10% DMSO to a concentration of 10mg/ml. Mice were administered 100mg/kg/day of the Favipiravir solution by i.p. injection, daily for up to 8 days.

#### IFN blocking antibody treatment

Mice were treated initially with 1mg and then 500μg every 3 days of BioXcell InVivoMAb anti-mouse IFNγ (cat: BE0055, clone: XMG1.2), InVivoMAb anti-mouse IFNAR-1 (cat: BE0241, clone: MAR1-5A3) or InVivoMAb polyclonal Armenian Hamster IgG isotype control (cat: BE0091) for up to 2 weeks during indicated treatment windows.

#### Tetramer generation and conjugation

A decoy tetramer was used to gate out cells binding to irrelevant tetramer components. The decoy tetramer consisted of DyLight 650 (DL650) conjugated to SA-PE loaded with an irrelevant protein, specifically biotinylated MHC-peptide (HLA-A2-M1).

#### Biotinylated protein production

Recombinant protein was expressed and produced by *Escherichia coli* (*E.coli*; BL21 strain). Transfected *E. coli* bacteria were selected in kanamycin (pET30 plasmid) or chloramphenicol containing medium overnight at 37°C, and then passaged to cultures at 1:50 and incubated at 37°C to reach an OD600nm of ∼0.6. Isopropyl β-D-1-thiogalactopyranoside (IPTG; Sigma-Aldrich) was added to cultures prior to further incubation at 37°C. Cells were centrifuged, lysed with lysozyme (10 mg per litre of expression) in the presence of DNAse. Supernatant was separated via centrifugation. NP peptidase was purified either by passing supernatant over a HisTrap Fast Flow column (Cytiva) or using a His-Bind purificatioin kit (EMD Chemicals). Purified protein was separated from *E. coli* proteins using a size exclusion column (Cytiva or GE Healthcare). For biotinylation of the protein, the recombinant protein was incubated with BirA enzyme and reaction buffer (50 mM Bicine pH 8.3, 10mM ATP, 10 mM magnesium acetate (MgOAc), and 50μM d-Biotin) at 4°C overnight. 1mg of protein per 10μg of enzyme was used and biotinylation efficiency was quantified using gel electrophoresis.

#### Tetramer conjugation

The Decoy tetramer was prepared by conjugating the core fluorochrome SA-PE to DL650 (Sigma-Aldrich) according to the manufacturer’s protocol for 60 minutes at room temperature. The free DL650 was removed by centrifugation in a 100-kD molecular weight cut off Amicon Ultra filter (Millipore). The SA-PE*DL650 complex concentration was calculated by measuring the absorbance of PE at 566nm using a NanoDrop ND-1000 spectrophotometer (Thermo Fisher Scientific). The SA-PE*DL650 complex was then incubated with 10-fold molar excess of biotinylated HLA-A2-M1 for 30 minutes at room temperature. The solution was diluted to 1μM based on the absorbance of PE at 566 nm (divided by the extinction coefficient = 1.96 cm−1μM−1). The LCMV-specific antigen tetramer was prepared by conjugating the biotinylated LCMV-NP to SA-PE (Prozyme PJRS25) at a concentration ratio of 4:1 respectively. Following addition of SA-PE to the biotinylated LCMV-NP, the mixture was first incubated in the dark at room temperature for 3 hours, then at 4°C overnight. The tetramer fraction was centrifuged in a 100-kD molecular weight cutoff Amicon Ultra filter. The concentration of tetramer was calculated by measuring the absorbance of PE at 566 nM (divided by the extinction coefficient = 1.96 cm−1μM−1).

#### Flow cytometric analysis and Fluorescence-activated cell sorting (FACS)

Single cell suspensions were isolated from each spleen. To collect peripheral blood mononuclear cells (PBMCs), needles were coated with heparin (StemCell) prior to cardiac puncture. Blood was layered onto an equal volume of Histopaque 1077 (Sigma-Aldrich) and centrifuged at 400g for 30 minutes at room temperature without brakes.

*Antigen-specific B cell enrichment:* The EasySep Mouse PE positive selection kit II (Stemcell) was used to enrich antigen-specific B cells using the manufacturer’s protocol. Enriched cells were filtered and resuspended in 1ml PBS 2% FCS. 2-5 x 10^6^ cells were added to FACS tubes and centrifuged (1500 RPM, 5 mins, 4°C). Cells were first stained with viability dye, Fixable Viability Stain 700 (FVS700 - BD Biosciences 564997) or FVS780 (BD Biosciences 565388). Cells were washed and resuspended in the relevant surface antibody staining cocktail. Following antibody staining, LCMV-infected samples were fixed using BD Cytofix according to the manufacturer’s protocol. Samples were analysed using a BD-LSRFortessa X-20 flow cytometer with FACSDiva software (BD Biosciences). FCS files were analysed using FlowJo software (FlowJo, LLC).

*Cell cycle analysis:* 5-Bromo-2’-deoxyuridine (BrdU, Sigma Aldrich) staining was performed as described previously^94^.

#### Assessment of viremia

The right lobe of the liver was collected from experimental mice. Livers were weighed and cDMEM was added to obtain 500 mg/ml of liver suspension. One stainless steel bead (Qiagen) was added per sample and liver tissues were homogenized using the TissueLyser LT (Qiagen) for 5 minutes at 50 oscillations/second. The beads were removed, and samples were centrifuged. The supernatant was collected for each sample and 10-fold serial dilutions were added to a 96-well culture plate (Falcon). A plaque forming assay was carried out as described previously^4^.

#### scRNA-seq

*Multiplexing samples with hashtag antibodies:* 100,000 cells were sort-purified per mouse. Cells were resuspended in washing buffer (PBS 0.04% BSA) and incubated with mouse Fc blockers (FcX, Biolegend). Each sample was then incubated with either TotalSeq anti-mouse Hashtag Antibody 1 (Biolegend #155861) or 2 (Biolegend #155863) for 20 minutes at 4°C. Cells were washed 3 times with 1ml of washing buffer to remove unbound antibodies. Following the final wash, LCMV-WE and LCMV-Docile samples were pooled at equal ratios in 1ml washing buffer. Pooled samples were then resuspended in PBS 0.04% BSA 0.2 U/ml RNAse inhibitor.

*Droplet-based scRNA-seq:* Reverse transcription, cDNA amplification and library preparation for single cell gene expression and V(D)J clonotypes were performed based on the manufacturer’s protocol using the Chromium Single Cell 5′ Library & Gel Bead Kit v3 (10X Genomics, Pleasanton, CA). Libraries were pooled at equimolar ratios and gene expression libraries were sequenced on the Illumina NovaSeq 6000 System, while HTO (hashtag oligo) and V(D)J (BCR) libraries were sequenced on Illumina NextSeq 500.

*Raw data processing:* Cellranger (v3.1.0) was used to align the raw scRNAseq data to the mouse reference genome (mm10-3.0.0). Cellranger count was run under default parameters. Cellranger VDJ (v3.1.0) was run under default parameters on the VDJ libraries to assemble and annotate VDJ sequences using the VDJ compatible reference (cellranger-vdj-GRCm38-alts-ensembl-3.1.0). To process the CITE-seq data, CITE-seq count (v1.4.3) was run with the following parameters (‘-- sliding window -cbf 1 -cbl 16 -umif 17 -umil 26 -cells $(number_submitted_cells)‘). Each sample was processed individually with each pipeline.

*Analysis of scRNA-seq data:* The Seurat package (v3.1.4) was used with R (v3.6)^95^. Count matrices from the CITE-seq pipeline and cellranger count were loaded into R and Seurat was used to demultiplex the data with the HTODemux function^96, 97^. Cells that lacked a corresponding hashtag were discarded. The two samples were merged after demultiplexing. Cells were stringently filtered for a minimum number of 1000 UMI, at least 500 genes detected and less than 5% mitochondrial reads for a cell to be kept. All doublets and empty cells were filtered out of the data, and a total of 6565 cells were retained. The data was scaled, log normalised and the number of principle components (PC) to be used for dimensionality reduction was determined using an elbow-plot. The first 10 PCs were selected, though the inclusion of more PCs was examined and found to have no change on the UMAP generated. The FindClusters function was used to cluster the data and was run with several different resolutions before settling on a resolution of 0.4, all other parameters were set to default. For all differential tests, the Mann-Whitney U test was used, and only positive markers were returned.

*Ingenuity pathway analysis:* Genes from scRNAseq datasets with a log fold change (FC) ≥2 with a false discovery rate (FDR, adjusted P-value) of <0.05 were defined as differentially expressed genes (DEGs) and assessed using Ingenuity Pathway Analysis software (IPA, Ingenuity Systems, QIAGEN). DEG data were uploaded to IPA for core analysis and analysed with the global molecular network in the Ingenuity Pathways Knowledge Base (IPKB). IPA identified canonical pathways, diseases and functions and molecular networks enriched by differentially expressed genes (DEGs)^98^.

*Gene set enrichment analysis (GSEA):* Gene expression profiles were generated using Loupe Browser (v5.0 - 10X Genomics) from LCMV-WE and LCMV-Docile samples. Genes were ordered in a ranked list based on differential expression between conditions. Gene set enrichment analysis (GSEA) (v4.1.0 - Broad Institute) was conducted with Signal2Noise values to determine where a priori sets of genes were distributed throughout the ranked list^99^. Genes related to a previously published phenotypic distinction will distribute at the top or bottom of the ranked list. Gene matrix files were obtained from MSigDB (Molecular Signature Database)^100^. The number of permutations was 1000, and the permutation type was set to gene set with FDR q-value less than 0.05.

##### Processing of single cell (V(D)J) datasets

FASTQ files for Chromium Single Cell V(D)J libraries were processed with 10x Genomics Cell Ranger 6.0.2 using the mouse VDJ reference (refdata-cellranger-vdj-GRCm38-alts-ensembl-5.0.0). Contigs generated by Cell Ranger (filtered_contigs.fasta) were post-processed with stand-alone IgBLAST (version 1.14.0) [doi: https://doi.org/10.1093/nar/gkt382] against the mouse V, D and J germline datasets obtained from the IMGT Reference Directories [website: https://www.imgt.org/vquest/refseqh.html] (accessed 16-Jan-2020) to produce tab-delimited output files (--outfmt 19).

Cell Ranger and IgBLAST output were merged with a perl script to provide a single line summary of heavy and light chains for each cell barcode. Where multiple chains were present at a single locus (i.e. IGH or IGK/L) the chain with the highest UMI count was retained and the presence of an additional chain was noted. VDJ data was merged by cell barcodes with single cell gene expression (GE) analysis in R version 4.2.1^101^ using the tidyverse package^102^. R was used within RStudio IDE^103^ and additional packages for data manipulation, analysis and visualisation included ggsci (colour palettes)^104^ and ggpubr (figure panel layouts)^105^.

Analysis for immunoglobulin repertoire features was restricted to cells with paired heavy and light chains. V gene somatic hypermutation (SHM) was derived from the IgBLAST output using the v_identity field (100 – v_identity). Isotype usage was determined from the Cell Ranger c_gene field for IGH chains with all IgG subclasses (IGHG1, IGHG3, IGHG2B/C) combined to a single IgG group and IgM sequences were split into ‘mutated’ (> 0% SHM) and ‘unmutated’ (0% SHM).

#### scATAC-seq

*Droplet-based scATAC-seq:* 100,000 cells were sort-purified per mouse. Nuclei isolation, barcoding and library preparation were performed according to the manufacturer’s protocol using the Chromium Next GEM Single Cell ATAC Reagent Kit v1.1 (10X Genomics). Libraries were prepared using the MGIEasy v3 chemistry kit and sequenced using the MGITech MGISEQ2000RS sequencer (MGI Tech).

*Raw data processing:* The raw sequence data was processed using cellranger-atac-2.0.0 for each sample against mm10, then aggregated using cellranger with normalize=none.

*Analysis of scRNA-seq data:* The aggregated data was loaded into R (v4.1.0) and analysed using Seurat (v4.1.0)^106^, and Signac (v1.5.0)^107^ QC filtering for peaks and cells was performed using these criteria: peaks present in at least 10 cells, cells that contained at least 200 called peaks and at least 10,000 fragments in peak regions. The standard Signac processing was performed, using dimension reduction with the LSI technique - the first dimension captured sequencing depth variation so was omitted from clustering and UMAP projection. Clustering was computed across several resolutions, and a resolution setting of 0.4 was chosen to give a good, stable clustering of 10 clusters. Clusters identified to contain contaminating cells were filtered out leaving a total of 7 clusters. Differentially accessible regions were found using the FindMarkers function from Seurat using the logistic regression (lr) method, testing regions that had detected fragments in at least 5% of cells in either cluster. The called peaks were then annotated using ChIPseeker^108^ to label where they occurred relative to the genes in the mm10 reference. This annotation per-peak was then broken down into per-cluster, and per-condition. ChromVAR^53^ was used to calculate significant differential motif activity between clusters. The motifs used were from the JASPAR 2020 database of transcription factor motifs^109^. First motif activity was calculated for each motif in each cell using the runChromVar function, then differential motif activity was determined using the FindMarkers function for a specific cluster compared to all other cells. Peaks found to be differentially accessible using the method described above were then selected to test for motif enrichment. Motif enrichment was tested using the findMotifsGenome method from HOMER^52^ which tests motifs found in the peaks against a background selected from the genome to balance for GC bias. The scRNA-seq cluster labels were transferred over to the scATAC-seq data using a standard approach. The “Gene Activity” for each gene, and each cell in the ATAC data was calculated based on the called peaks falling within genes. The FindTransferAnchors function was then used to identify mutual nearest neighbours in a projected lower dimensional space between the two datasets, then these anchor cells were used to transfer the scRNA-seq cluster labels to the nearby cells in the scATAC-seq dataset.

#### Bulk ATAC-seq

Approximately 15,000 cells were pelleted and washed with PBS. One-step permeabilization and tagmentation method was used by resuspending the cell pellets in Digitonin (EZSolution), Tween-20 (Sigma) buffer. High-molecular-weight tagmented DNA was then removed by incubating samples with a 0.7X ratio of Agencourt AMPureXP beads (Invitrogen). The same process was applied using a 1.2X ratio of beads to exclude low-molecular-weight DNA. Purified DNA was then pre-amplified with indexed primers and HiFi HotStart Polymerase Ready Mix (KAPA Biosystems). After estimation of additional PCR cycles required^110^, the genomic library was fully amplified using the SYBR qPCR Master Mix with primers included in the Illumina Library Quantification kit for Bio-Rad iCycler (KAPA Biosystems). Libraries were sequenced on the Illumina NextSeq 2000 using 60-nt paired-end Illumina chemistry. Analysis was performed as described previously^4^.

### Statistical analysis

Experimental data are presented as mean ± standard error of the mean (SEM) with statistical analysis performed using GraphPad Prism 8 (GraphPad Software). When comparing groups, between-group differences were analysed using the Mann-Whitney non-parametric two-tailed test with a 95% confidence interval.

## DATA AND CODE AVAILABILITY

The sequencing data will be deposited will be made available to the public upon publication.

---

**Figure S1.**
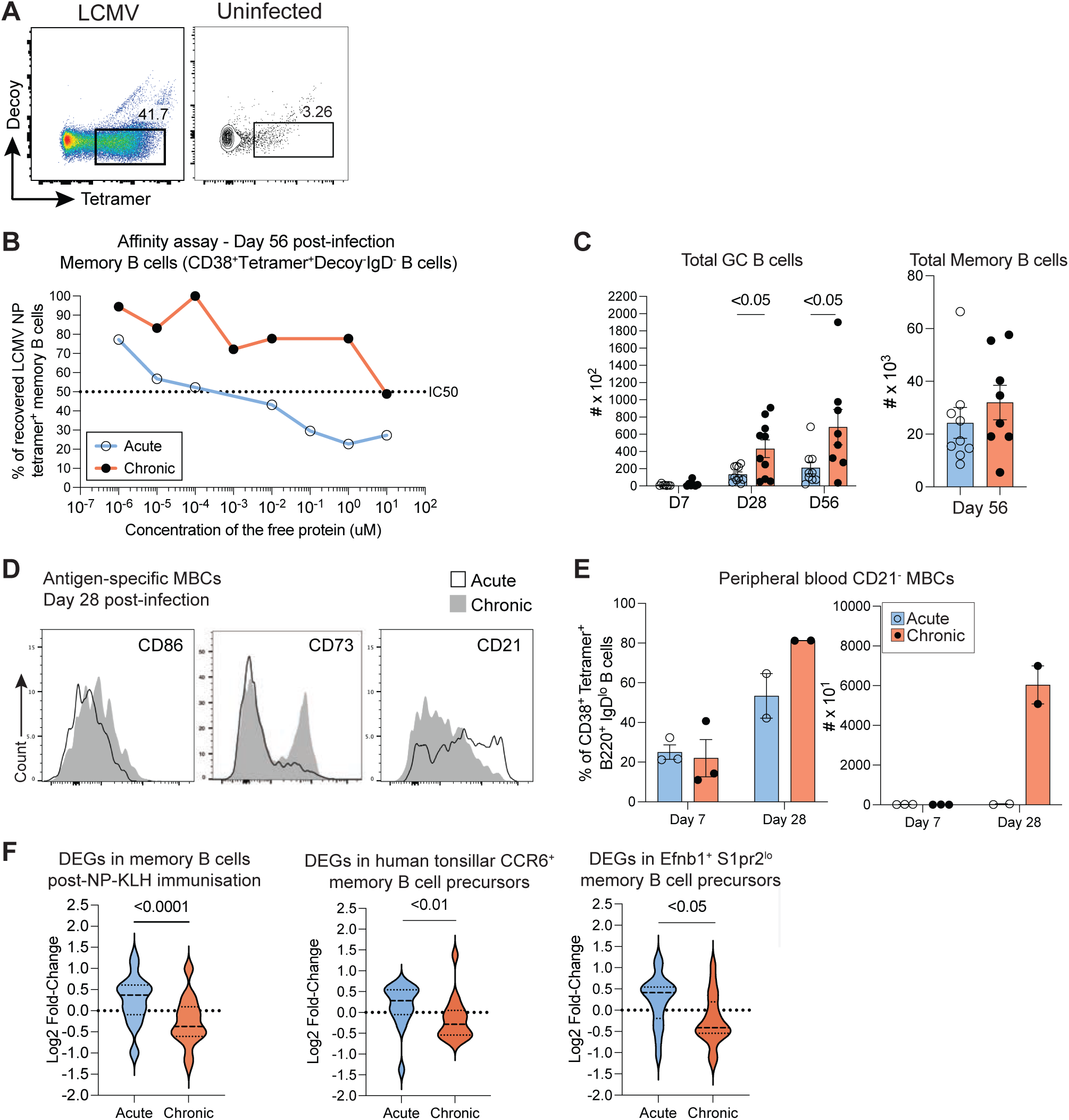
Memory B cell affinity for LCMV NP antigen is higher in acute versus chronic LCMV infection at all timepoints assessed. (A) Representative flow cytometry plots of Tetramer^+^ antigen-specific B cells (B220^+^IgD^−^ Tetramer^+^Decoy^−^) at 14 days post-LCMV infection, compared to an uninfected control. (B) Graph representing affinity of the B cells for the tetramer from LCMV-WE or LCMV-Docile-infected mice, with the concentration of monomeric antigen required to prevent 50% tetramer binding represented as IC50. (C) Total number of splenic GC B cells (B220^+^IgD^−^Tetramer^+^Decoy^−^CD38^−^CD95^hi^) at day 7 (WE: n = 7, Docile: n = 7), day 28 (WE: n = 10, Docile: n = 10), or day 56 (WE: n = 10, Docile: n = 9) post-LCMV-WE or LCMV-Docile infection; splenic memory B cells (B220^+^IgD^−^ Tetramer^+^Decoy^−^CD38^−^) at d56 (WE: n = 9, Docile: n = 8). Combined data from 8 independent experiments. Data represent mean ± SEM. (D) Representative histograms of CD86, CD73 and CD21 gated from enriched splenic memory B cells (B220^+^IgD^−^Tetramer^+^Decoy^−^CD38^+^) from mice infected with LCMV-WE or LCMV-Docile at day 28 post-infection. (E) Flow cytometric analyses of CD21^−^ frequency within the peripheral blood memory B cell population and total number at day 7 (WE: n = 3, Docile: n = 3) or day 28 (WE: n = 2, Docile: n = 2; each replicate is pooled from two individual donors) post-LCMV-WE or LCMV-Docile infection. Combined data from 2 independent experiments. Data represent mean ± SEM. (F) Violin plots showing log2 fold-change expression of genes associated with memory B cells post-NP-KLH immunisation, Ephrin-B1^+^ memory precursors and CCR6^+^ memory precursors in memory B cells (*Cd38*^+^*Ighd*^−^ non-GC) in LCMV-WE or LCMV-Docile-infected mice. Related to Figure 1.

**Figure S2.**
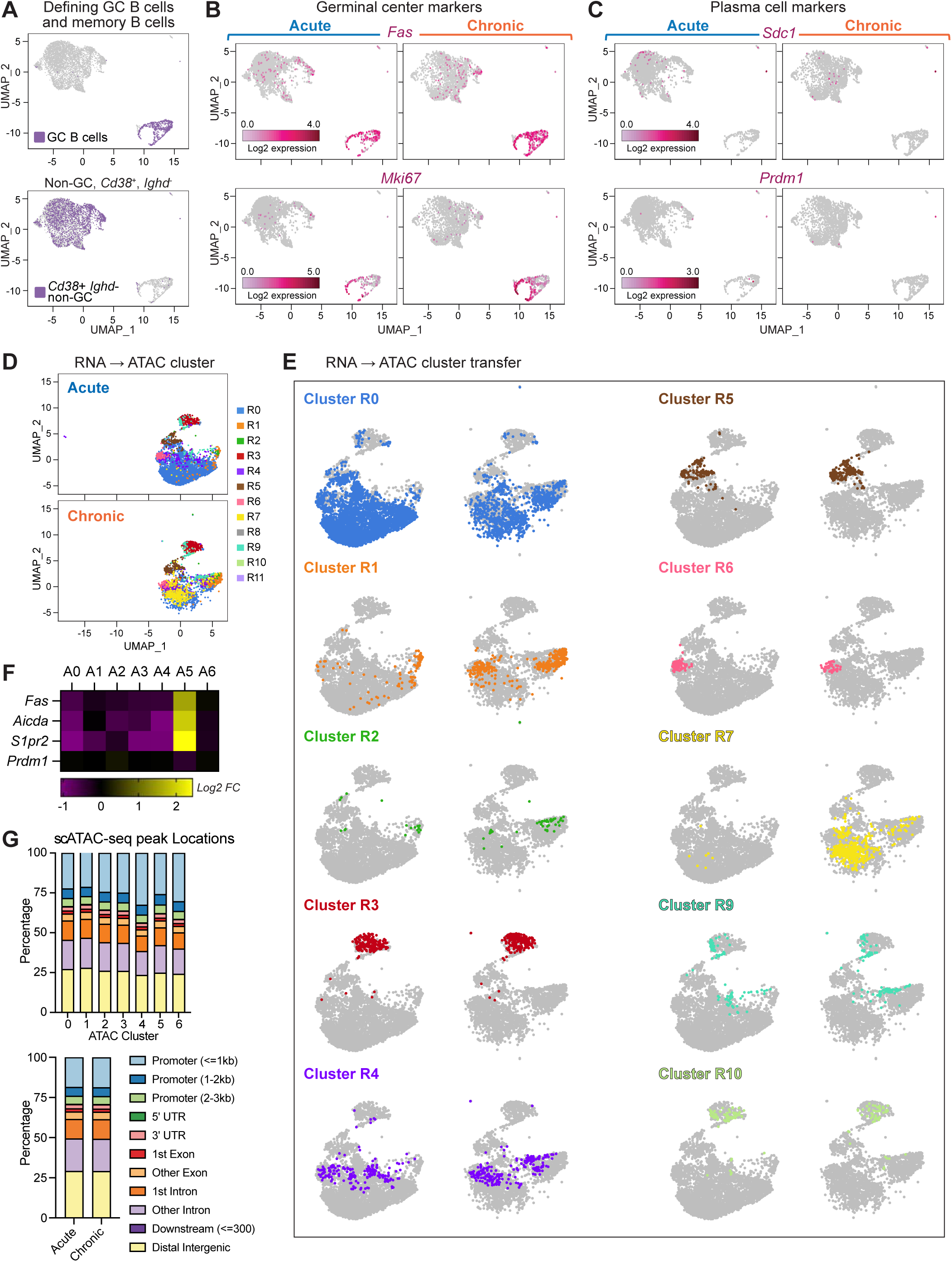
Clusters expanded in chronic LCMV infection within the single cell RNA-seq dataset corresponded to those expanded in the single cell ATAC-seq dataset. (A) UMAP of single cell RNA seq data, with GC B cells (top) and memory B cells (*Cd38*^+^ *Ighd*-non-GC cells) (bottom) shown as purple dots. (B and C) UMAPs of gene expression of (B) GC markers (*Fas* and *Mki67*) and (C) plasma cell markers (*Sdc1* and *Prdm1*) split by condition (LCMV-WE: left, LCMV-Docile: right), with positive expression represented by pink dots. (D and E) Visualization of label transfer. scATAC-seq UMAP plots split by infection type, with scRNAseq cluster annotations overlayed. Shown as (D) combined clusters or (E) each cluster highlighted. (F) Graph representing the number of cut sites per cell that fall within promoter and distal peaks near GC-associated genes, split by cluster. (G) Percentage of peak locations, split by condition and by cluster. Related to Figure 2.

**Figure S3.**
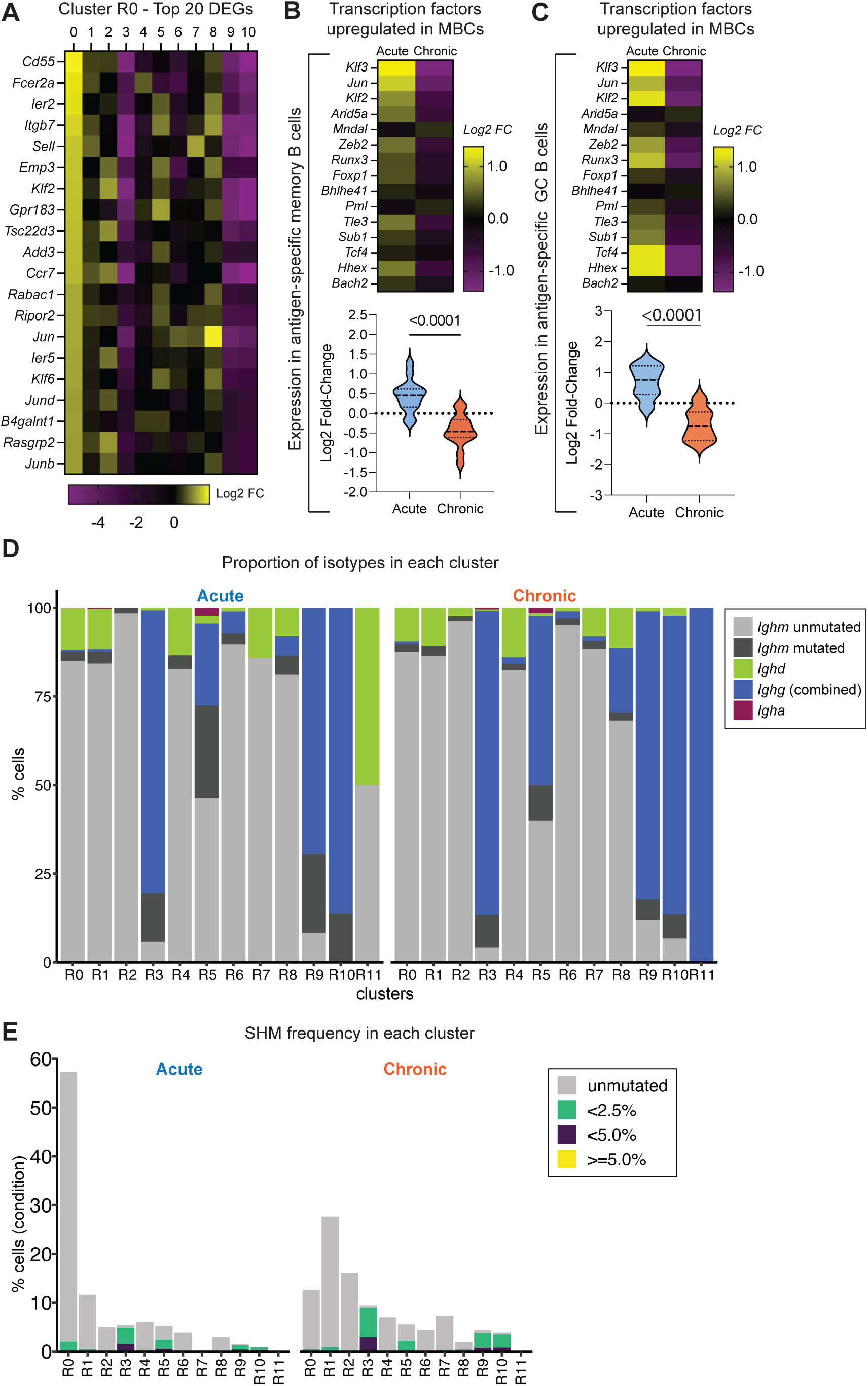
Gene sets associated with conventional memory B cell formation were increased in memory B cells in acute versus chronic LCMV infection. (A) Heatmap of the top 20 DEGs in cluster R0 compared to all other clusters (excluding Ig genes). (B and C) Heatmaps and violin plots representing log2 fold-change in expression of memory B cell-associated transcription factor genes in (B) antigen-specific GC B cells or (C) memory B cells (*Cd38*^+^*Ighd*^−^ non-GC) in LCMV-WE or LCMV-Docile infected mice. (D) scVDJ-seq: Graph representing the percentage of each antibody isotype within each cluster, split by infection type. (E) scVDJ-seq: Graph representing mutation frequency per cluster, split by infection type. Related to Figure 2.

**Figure S4.**
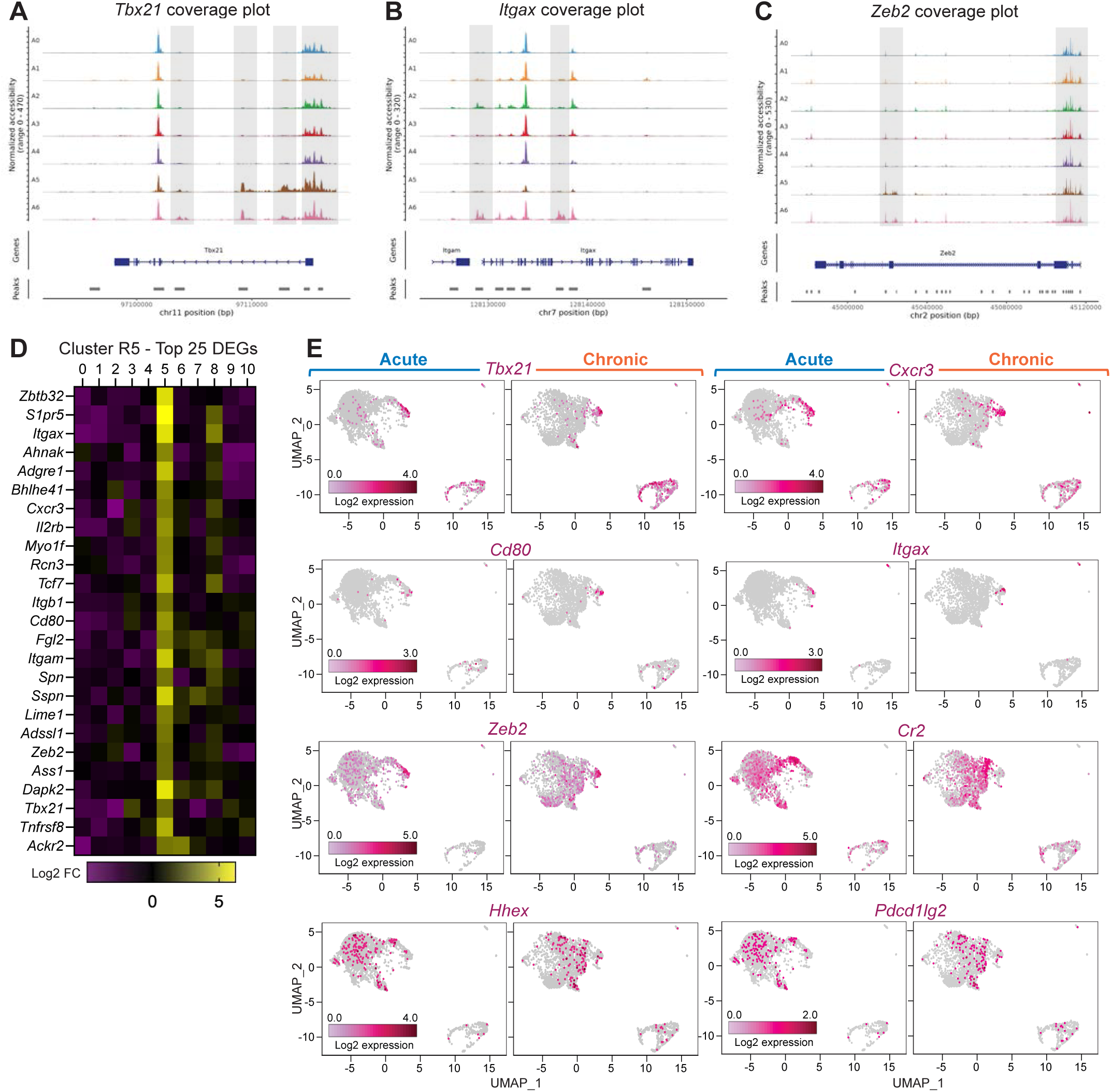
Memory B cell subsets expanded in chronic LCMV infection were distinct from previously described T-bet-associated memory B cells. (A, B and C) Coverage plots showing chromatin accessibility at the (A) *Tbx21* gene region, (B) *Itgax* gene region, or (C) *Zeb2* gene region, split by ATAC cluster. (D) Heatmap of the top 25 DEGs in cluster R5, compared to all other clusters. (E) UMAPs of gene expression of *Tbx21*, *Cxcr3*, *Cd80*, *Itgax*, *Zeb2*, *Cr2*, *Hhex* and *Pdcd1lg2* split by condition (LCMV-WE: left, LCMV-Docile: right), with positive expression represented by pink dots. Related to Figure 2.

**Figure S5.**
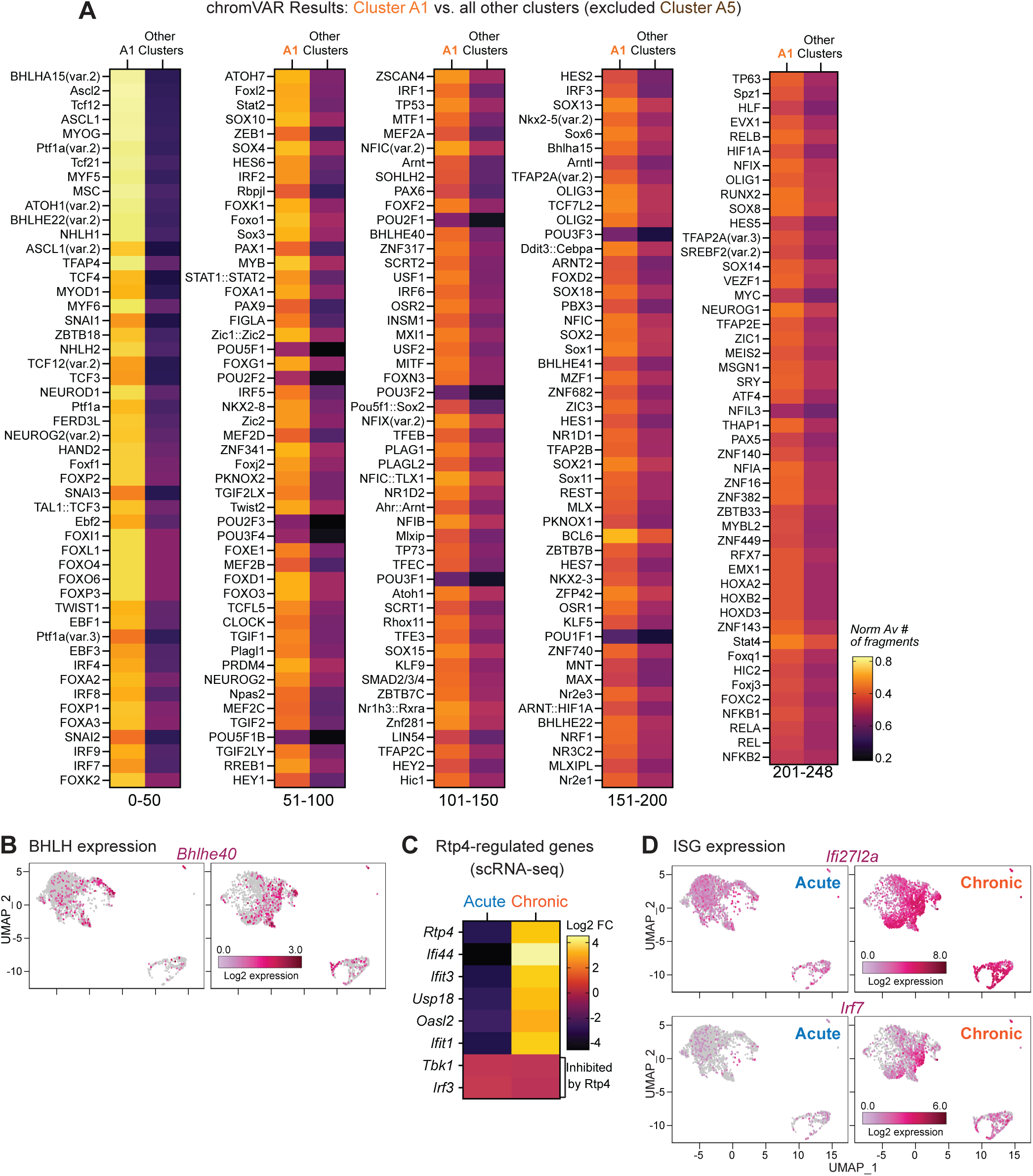
Epigenetic signatures in memory B cells in chronic LCMV infection showed enrichment of transcription factor motifs linked to regulation of B cell function and differentiation. (A) Heatmap of all (248) DARs in cluster A1 compared to all other clusters (excluding cluster A5). (B) UMAPs of gene expression of *Bhlhe40* split by condition (LCMV-WE: left, LCMV-Docile: right), with positive expression represented by pink dots. (C) Heatmap of gene expression of Rtp4-associated genes in memory B cells (*Cd38*^+^*Ighd*^−^ non-GC) in LCMV-WE vs. LCMV-Docile infection (generated using single cell RNA-seq data). (D) UMAPs of gene expression of *Ifi27l2a* and *Irf7* split by condition (LCMV-WE: left, LCMV-Docile: right), with positive expression represented by pink dots. Related to Figure 3.

**Figure S6.**
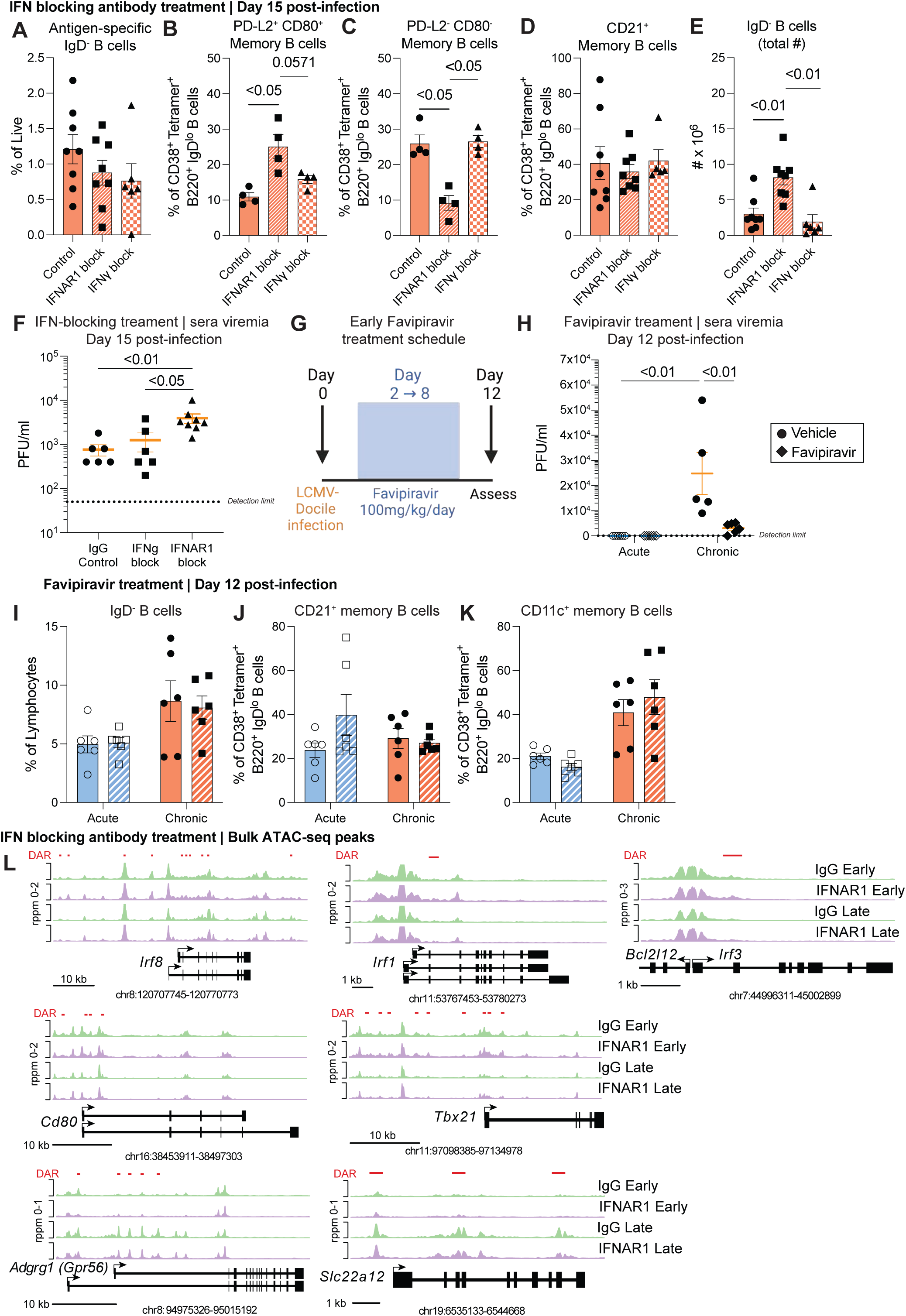
Type I IFN blocking antibody early in infection, but not modulation of viral load, alters memory B cell phenotype. (A) Flow cytometric analyses of antigen-specific B cells (B220^+^ IgD^−^ Tetramer^+^ Decoy^−^) in mice treated with anti-IFNAR1 (MAR1-5A3) (n = 8), anti-IFNγ (XMG1.2) (n = 5) or IgG control (n = 8) at day 15 post-infection. Combined data from 2 independent experiments. Data represents mean ± SEM. (B, C and D) Flow cytometric analyses of (B) PD-L2^+^CD80^+^ frequency within memory B cells, (C) PD-L2^−^CD80^−^ frequency within memory B cells and (D) CD21^+^ frequency within memory B cells in mice treated with anti-IFNAR1 (MAR1-5A3) (n = 4), anti-IFNγ (XMG1.2) (n = 3) or IgG control (n = 4) at day 15 post-infection. Combined data from 2 independent experiments. Data represents mean ± SEM. (E) Flow cytometric analyses of activated B cells (B220^+^IgD^−^) in mice treated with anti-IFNAR1 (MAR1-5A3) (n = 8), anti-IFNγ (XMG1.2) (n = 5) or IgG control (n = 8) at day 15 post-infection. Combined data from 2 independent experiments. Data represents mean ± SEM. (F) Sera LCMV viremia (PFU/ml) as detected via plaque forming assay in mice treated with anti-IFNAR1 (MAR1-5A3) (n = 6), anti-IFNγ (XMG1.2) (n = 6) or IgG control (n = 8) at day 15 post-infection. Combined data from 2 independent experiments. Data represents mean ± SEM. (G) Schematic of Favipiravir treatment schedule and day 12-post infection timepoint. (H) Sera LCMV viremia (PFU/ml) as detected via plaque forming assay in mice treated with Favipiravir (T-705) (WE: n = 6, Docile: n = 6), or Vehicle control (WE: n = 6, Docile: n = 5) at day 12 post-infection. Combined data from 2 independent experiments. Data represents mean ± SEM. (I, J, and K) Flow cytometric analyses at day 12 post-infection of (I) activated B cells (B220^+^ IgD^−^), (J) CD21^+^ frequency within memory B cells (B220^+^IgD^−^Tetramer^+^Decoy^−^CD38^+^) and (K) CD11c^+^ frequency within memory B cells in mice treated with Favipiravir (T-705) (WE: n = 2, Docile: n = 2), or Vehicle control (WE: n = 6, Docile: n = 6). Combined data from 2 independent experiments. Data represents mean ± SEM. (L) Coverage plots showing chromatin accessibility at the *Irf8, Irf1, Irf3, Cd80, Tbx21, Adgrg1(Gpr56)* and *Slc22a12* gene regions, split by treatment group. Related to Figure 4, Figure 5 and Figure 6.

